# *In vivo*-compatible spatial multi-omics via hydrogen peroxide-independent APEX2 labeling

**DOI:** 10.64898/2026.03.13.711744

**Authors:** Boyi Chen, Hongyang Guo, Zhijian Yan, Wenjie Lu, Chun Li, Shimin Xu, Yanling Zhang, Haodong Guo, Shaopeng Sun, Xuege Sun, Shangxun Zhao, Qiaoling Shangguan, Yidong Chen, Lei Lu, Zhaofa Wu, Ying Chen, Wei Qin

## Abstract

Proximity labeling (PL) technologies like APEX2 have transformed spatial multi-omics in live cells, but their long-standing dependence on hydrogen peroxide (H₂O₂) disrupts redox signaling and prevents use in live animals. Here we introduce H₂O₂-independent APEX2 (Hi-APEX), which uses a clickable tetrazine-phenol probe, requiring no enzyme engineering. We show that APEX2 directly catalyzes TP radical formation without H₂O₂ via a mechanism requiring the probe’s tetrazine group and a key histidine residue. We benchmarked Hi-APEX-based spatial multi-omics by mapping the mitochondrial matrix and dynamic secretomes. Hi-APEX significantly outperforms traditional APEX in capturing redox-sensitive processes such as stress response and ferroptosis, enabling discovering authentic stress granule components and protein interaction networks for mitochondria-localized GPx4. One mGPx4 interactor TRMT61B—known to regulate mitochondrial m¹A modifications—promotes ferroptosis. Crucially, Hi-APEX achieves full *in vivo* compatibility, enabling direct PL in tumor xenografts and hippocampal neurons, thereby expanding PL-based spatial multi-omics from cellular systems to living organisms.

---

Cellular function in mammals relies on intricate compartmentalization, where biomolecules organize into specialized assemblies that execute compartment-specific functions^1^. To decode this architectural complexity, researchers have developed spatial multi-omics approaches that map biomolecules across scales—from tissue-level organization down to dynamic subcellular microdomains^2,3^. While traditional fractionation-based methods have provided valuable insights, they often lack the resolution to capture transient molecular interactions within specialized cellular niches^4^.

Among the most powerful tools to decode molecular architectures are proximity labeling (PL) techniques^5^, particularly those utilizing engineered peroxidases like APEX2^6^. These enzymes generate reactive radicals that covalently tag nearby biomolecules with nanometer-scale precision when activated by H₂O₂. APEX2’s exceptional temporal resolution (<1 minute) has enabled groundbreaking studies of elusive cellular compartments and dynamic processes, bridging proteomic and transcriptomic mapping at unprecedented resolution^7–10^.

Despite this potential, peroxidase-based PL methods, including APEX2 and horseradish peroxidase (HRP), remain constrained by the inherent toxicity of H₂O₂, restricting applications to cultured cells or *ex vivo* tissues and precluding live animal studies. Even in transgenic models, labeling requires tissue dissection and immersion in H₂O₂ buffers, bypassing true *in vivo* operation^11,12^. Moreover, H₂O₂ exposure disrupts redox homeostasis, confounding studies of redox-sensitive pathways. Recent attempts to circumvent exogenous H₂O₂, such as fusing APEX2 with H₂O₂-generating enzymes, also risk altering redox balance and introducing light-dependent cytotoxicity^13^. Alternative PL strategies face their own limitations: promiscuous biotin ligases suffer from high background noise and lack RNA-labeling capability^14^; tyrosinase- and laccase-based methods grapple with copper toxicity and spatial restriction^15–17;^ and photocatalyst-dependent systems, while enabling simultaneous protein/RNA labeling, are hampered by poor tissue penetration of visible light^18–20^.

To overcome these challenges, we developed Hi-APEX, a novel PL platform that utilizes tetrazine-phenol (TP) as an H₂O₂-independent substrate for APEX2 (**Fig. 1a**). This system enables efficient protein labeling without exogenous H₂O₂, instead leveraging endogenous peroxide levels while maintaining robust labeling efficiency. The tetrazine moiety of labeled proteins rapidly conjugates with TCO-biotin via a 10-minute inverse electron-demand Diels-Alder reaction^21^, facilitating straightforward streptavidin-based enrichment and multi-omic profiling. We have validated Hi-APEX across multiple biological contexts, demonstrating its unique capability to resolve dynamic processes particularly vulnerable to H₂O₂ perturbation in living cells and animals. As TP is commercially available and the IEDDA workflow is streamlined, Hi-APEX can be seamlessly integrated into existing APEX2 workflows without genetic modification, offering broad accessibility to the research community.

**Fig. 1.**
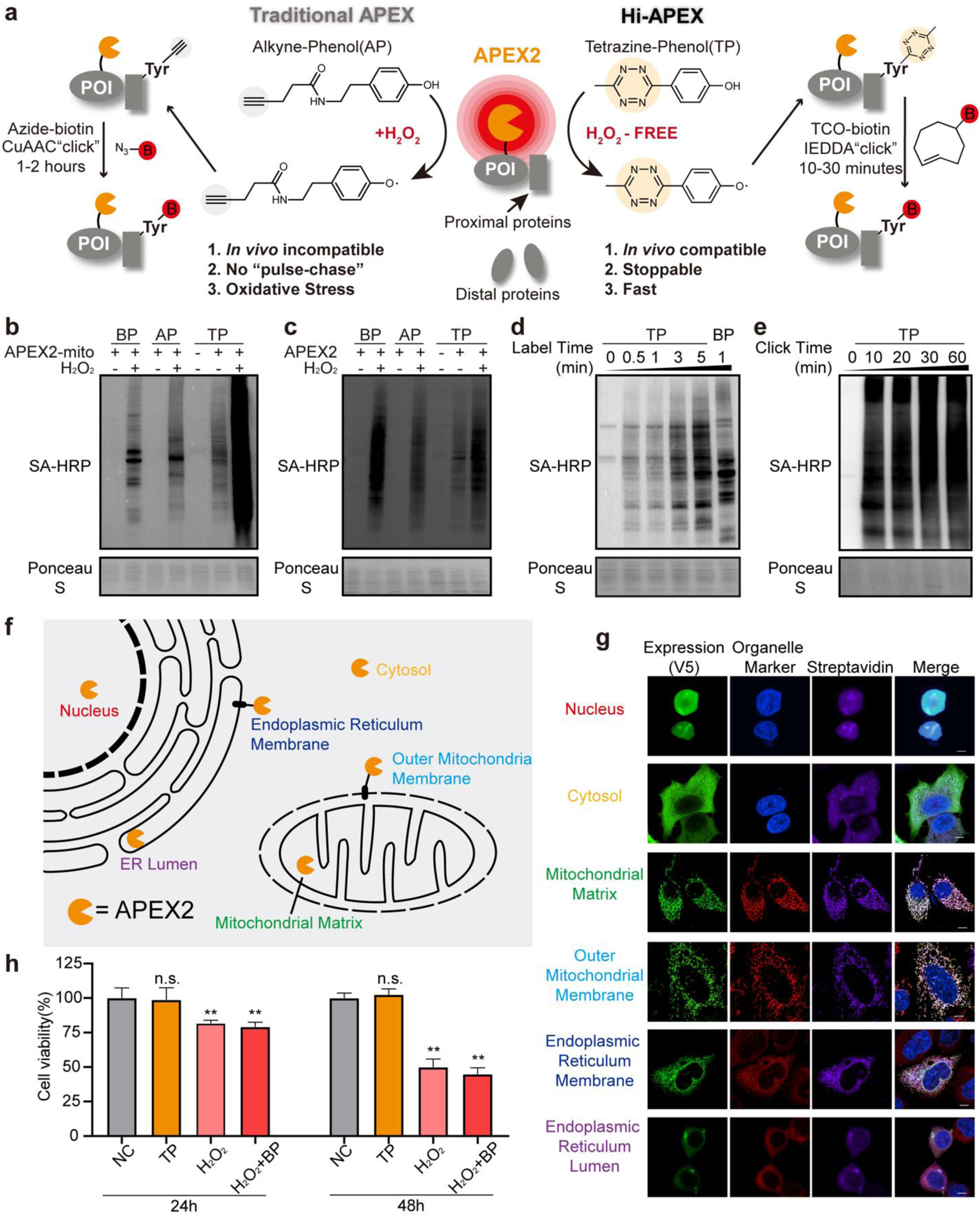
Discovery of tetrazine-phenol as an H₂O₂-independent substrate for APEX2 labeling. (a) Schematic comparing Hi-APEX with conventional APEX2 labeling. Conventional APEX2 requires a 1-minute pulse of exogenous H₂O₂, and labeling with alkyne-phenol (AP) necessitates a subsequent 1-hour copper-catalyzed azide-alkyne cycloaddition (CuAAC) for detection. In contrast, Hi-APEX labeling with tetrazine-phenol (TP) operates without exogenous H₂O₂, and detection is achieved via a 10-minute inverse electron-demand Diels-Alder (IEDDA) reaction. (b) Assessment of mitochondria-targeted APEX2 labeling in living HEK293T cells using biotin-phenol (BP), AP, and TP. Cells expressing APEX2-mito were treated with 500 µM BP for 30 minutes, 50 µM AP for 30 minutes, or 500 µM TP for 5 minutes, followed by a 1-minute pulse of either vehicle or 1 mM H₂O₂. After lysis, AP-labeled samples underwent CuAAC with azide-biotin, while TP-labeled samples were subjected to an IEDDA reaction with TCO-biotin. Biotin incorporation was analyzed by streptavidin blotting. (c) Evaluation of APEX2 labeling in HEK293T cell lysates. Lysates were incubated with 2 µM recombinant APEX2 and 500 µM of BP for 30 minutes, 500 µM of AP for 30 minutes or 500 µM of TP for 5 minutes, followed by a 1-minute vehicle or 1 mM H₂O₂ treatment. (d) Time course of APEX2 labeling with TP. HEK293T cells expressing APEX2-mito were treated with 500 µM TP for the indicated durations. (e) Optimization of the IEDDA reaction time. HEK293T cells expressing APEX2-mito were labeled with 500 µM TP for 5 minutes. Lysates were then incubated with 20 µM TCO-biotin for the specified times to conjugate the label. (f) Schematic of APEX2 targeted to distinct subcellular locales: cytosol, nucleus, mitochondrial matrix, outer mitochondrial membrane (OMM), endoplasmic reticulum membrane (ERM), and ER lumen. (g) Confocal microscopy images of TP-based APEX2 labeling. APEX2 was visualized with an anti-V5 antibody, and biotin labeling from the TP/TCO-biotin reaction was detected with streptavidin. Scale bars, 10 µm. (h) Cell viability assay of APEX2-mito cells 24 and 48 hours after labeling with either H₂O₂-free TP or H₂O₂-dependent BP. \*\**p* < 0.01; n.s., not significant. Data represented as mean ± SD.

## Discovery of Tetrazine-Phenol as an H₂O₂-Independent APEX2 Substrate

During prior development of non-biotin substrates for TransitID, we observed that alkyne-phenol (AP) provided more efficient H₂O₂-dependent labeling than biotin-phenol (BP), which we attributed to its superior membrane permeability^22^. Building on this, we unexpectedly discovered that a tetrazine-phenol (TP) derivative exhibited even stronger labeling—an enhancement not explained by permeability alone, as TP also showed higher reactivity in cell lysates (**Fig. 1b, c**). Remarkably, TP produced robust APEX2-dependent labeling even without exogenous H₂O₂, achieving signals comparable to BP with standard 1 mM H₂O₂. This H₂O₂-independent activity was consistent across multiple cell lines and was abolished by sodium ascorbate, confirming a phenol radical–mediated mechanism (**Supplementary Fig. 1a, b**).

APEX2-dependent TP labeling is both rapid and concentration-dependent. In cells expressing mitochondrial matrix–targeted APEX2, a 5-minute TP treatment yielded stronger labeling than the standard 1-minute BP/H₂O₂ protocol, with detectable signal emerging within one minute (**Fig. 1d**). The subsequent IEDDA conjugation with TCO-biotin was complete within 10 minutes, underscoring the kinetic efficiency of this approach (**Fig. 1e**).

We next validated the spatial specificity of TP labeling across multiple subcellular compartments—including the cytosol, nucleus, ER lumen and membrane, mitochondrial matrix and outer membrane—and observed strong, H₂O₂-independent labeling in each (**Supplementary Fig. 1c**). Confocal imaging confirmed precise colocalization of TP-derived signal with both APEX2 and organelle markers (**Fig. 1g**). The method also proved effective for cell-surface labeling using HRP-fused constructs, further supporting its broad applicability (**Supplementary Fig. 1d**).

A key advantage of TP labeling is its low toxicity. Unlike conventional APEX2 labeling, which requires immediate quenching due to H₂O₂ cytotoxicity, a 5-minute TP treatment did not affect cell viability (**Supplementary Fig. 1e**). In pulse-chase experiments, cells labeled with TP proliferated normally over 48 hours, whereas BP/H₂O₂–treated cells showed marked growth suppression (**Fig. 1h**). Streptavidin blotting confirmed that labeling did not increase after TP removal, suggesting TP labeling is stoppable (**Supplementary Fig. 1f**). Based on these results, we designate this method Hi-APEX (H₂O₂-independent APEX), offering a non-toxic and readily controllable alternative for PL.

## Confirmation and Mechanistic Evaluation of H_2_O_2_-independent TP labeling *in vitro*

To definitively establish whether APEX2-mediated TP labeling operates independently of H₂O₂, we performed *in vitro* assays with purified recombinant APEX2 and bovine serum albumin (BSA). Strikingly, APEX2 catalyzed TP labeling of BSA without any exogenous H₂O₂, while parallel BP reactions showed no signal when H₂O₂ was omitted (**Fig. 2a**). This TP labeling was quenched by sodium ascorbate. When supplemented with 10 µM H₂O₂—approximating physiological intracellular levels^23^—TP labeling intensity increased substantially (**Fig. 2b**). Similar H₂O₂-independent labeling was observed with HRP (**Supplementary Fig. 2a**). This suggests that peroxidases can directly catalyze TP labeling in the absence of H₂O₂, with low concentrations of H₂O₂ further facilitating this reaction.

**Fig. 2.**
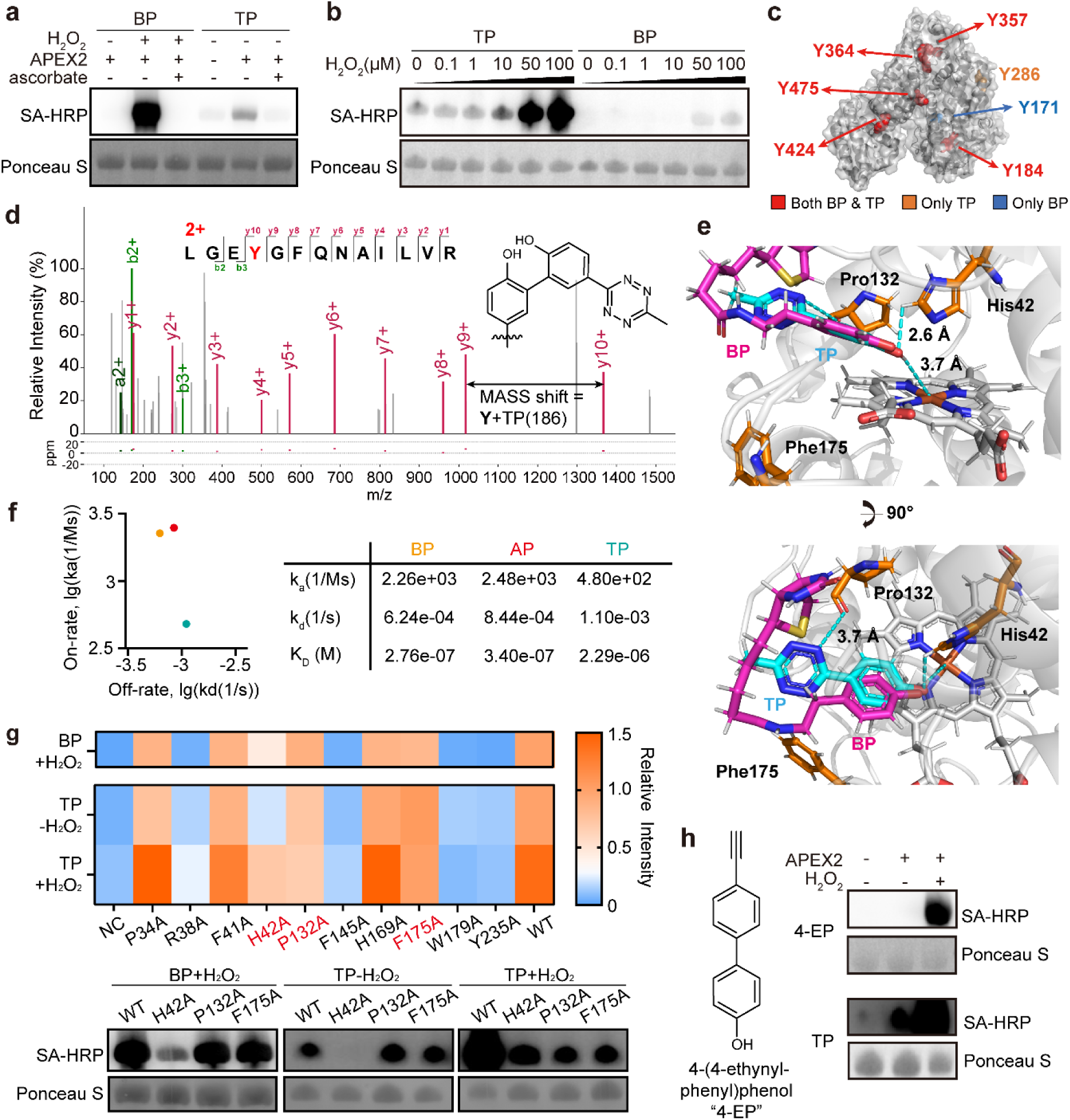
Mechanistic investigation of H₂O₂-independent TP labeling *in vitro*. (a) H₂O₂-independent TP labeling of BSA by recombinant APEX2. Reactions contained 2 µM APEX2 and 2 mg/mL BSA incubated with 500 µM TP for 30 min. For BP labeling, parallel reactions with 500 µM BP were activated with a 1-minute pulse of 1 mM H₂O₂. To probe the mechanism, samples were treated with 10 mM sodium ascorbate. TP-labeled proteins were subsequently conjugated via IEDDA with TCO-biotin, and all samples were analyzed by streptavidin blotting. (b) Effect of H₂O₂ concentration on *in vitro* APEX2 labeling. A mixture of 2 µM APEX2, 2 mg/mL BSA, and 500 µM TP or BP was treated with the indicated H₂O₂ concentrations. (c) Mapping of H₂O₂-independent TP labeling sites on BSA reveals overlap with conventional BP labeling sites. (d) Representative MS2 spectrum of a TP-modified BSA peptide. (e) Structural docking of TP and BP into the apo form of APEX2 (PDB: 9X9L), highlighting key residues His42, Pro132, and Phe175. (f) Binding affinities of APEX2 for different probes, as determined by surface plasmon resonance (SPR). (g) Effect of APEX2 point mutations on H₂O₂-independent TP labeling. The heat map shows relative labeling efficiency compared to wild-type (WT). Streptavidin blots for His42, Pro132, and Phe175 mutants are shown below. (h) *In vitro* APEX2 labeling with 4-(4-ethynylphenyl)phenol (4-EP). Reactions containing 2 µM APEX2 and 2 mg/mL BSA were incubated with 500 µM 4-EP and 1 mM H₂O₂.

Liquid chromatography-tandem mass spectrometry (LC-MS/MS) analysis of trypsin-digested, TP-labeled BSA revealed modifications on multiple surface-exposed tyrosine residues—overlapping with known APEX2 labeling sites for BP under H₂O₂ stimulation (**Fig. 2c**)^24^. The observed mass shift indicated a conjugation chemistry shared with BP modifications (**Fig. 2d and Supplementary Fig. 2b**).

Though we were unable to obtain a TP-APEX2 co-crystal structure, molecular docking using our solved apo-APEX2 structure positioned the phenol groups of TP, BP, and AP between the heme group and His42—the established site for phenoxyl radical generation (**Fig. 2e, Supplementary Fig. 2c, d**)^25,26^. The compact tetrazine moiety of TP was uniquely positioned near Pro132, where it likely forms a hydrogen bond. Surface plasmon resonance confirmed that TP binds APEX2 with weaker affinity and faster dissociation than BP or AP (**Fig. 2f, Supplementary Fig. 2e**), suggesting a more flexible interaction.

We selected ten residues in adjacent to TP and evaluated the impact of their mutation on TP labeling. The H42A mutant abolished H₂O₂-independent TP labeling while retaining residual H₂O₂-dependent activity (**Fig. 2g, Supplementary Fig. 2f**). Docking indicated altered TP positioning in this mutant (**Supplementary Fig. 2g**), underscoring His42’s essential role in H₂O₂-independent TP catalysis. Interestingly, P132A and F175A mutants efficiently catalyzed TP labeling without H₂O₂, yet showed no response to H₂O₂ supplementation (**Fig. 2g**), suggesting these residues specifically mediate H₂O₂-dependent TP activation.

The unique properties of TP that enable H₂O₂-independent labeling stem from its molecular architecture. TP contains a π-π conjugated system where the electron-withdrawing tetrazine group modulates the phenol’s electronic properties. To test if this specific moiety is essential, we used 4-(4-ethynylphenyl)phenol, a compound with a similar conjugated system. This analog showed the same APEX2 binding form as TP but failed to produce any H₂O₂-independent labeling (**Fig. 2h; Supplementary Fig. 2h**). These results underscore that the tetrazine group confers the unique catalytic ability for H₂O₂-independent labeling.

## Proteomic and transcriptomic benchmark of Hi-APEX

To benchmark Hi-APEX, we performed proteomic profiling in the mitochondrial matrix—a well-defined compartment for evaluating PL methods^7^. Following TP labeling, proteins were conjugated to TCO-biotin, enriched by streptavidin pulldown, and analyzed using data-independent acquisition (DIA)-based LC-MS/MS (**Fig. 3a**). We quantified 4270 proteins across three biological replicates with high reproducibility (**Supplementary Fig. 3a, Table S1**). Using mitochondrial matrix and cytosolic proteins as true and false positives, respectively, ROC analysis confirmed strong enrichment of mitochondrial proteins by Hi-APEX (**Fig. 3b**). Applying a cutoff maximizing the difference between TP and FP rates, Hi-APEX-enriched proteins (*p* < 0.05; **Fig. 3c**) showed strong enrichment for mitochondrial pathways and specifically localized to the matrix (**Fig. 3d, e**).

**Fig. 3.**
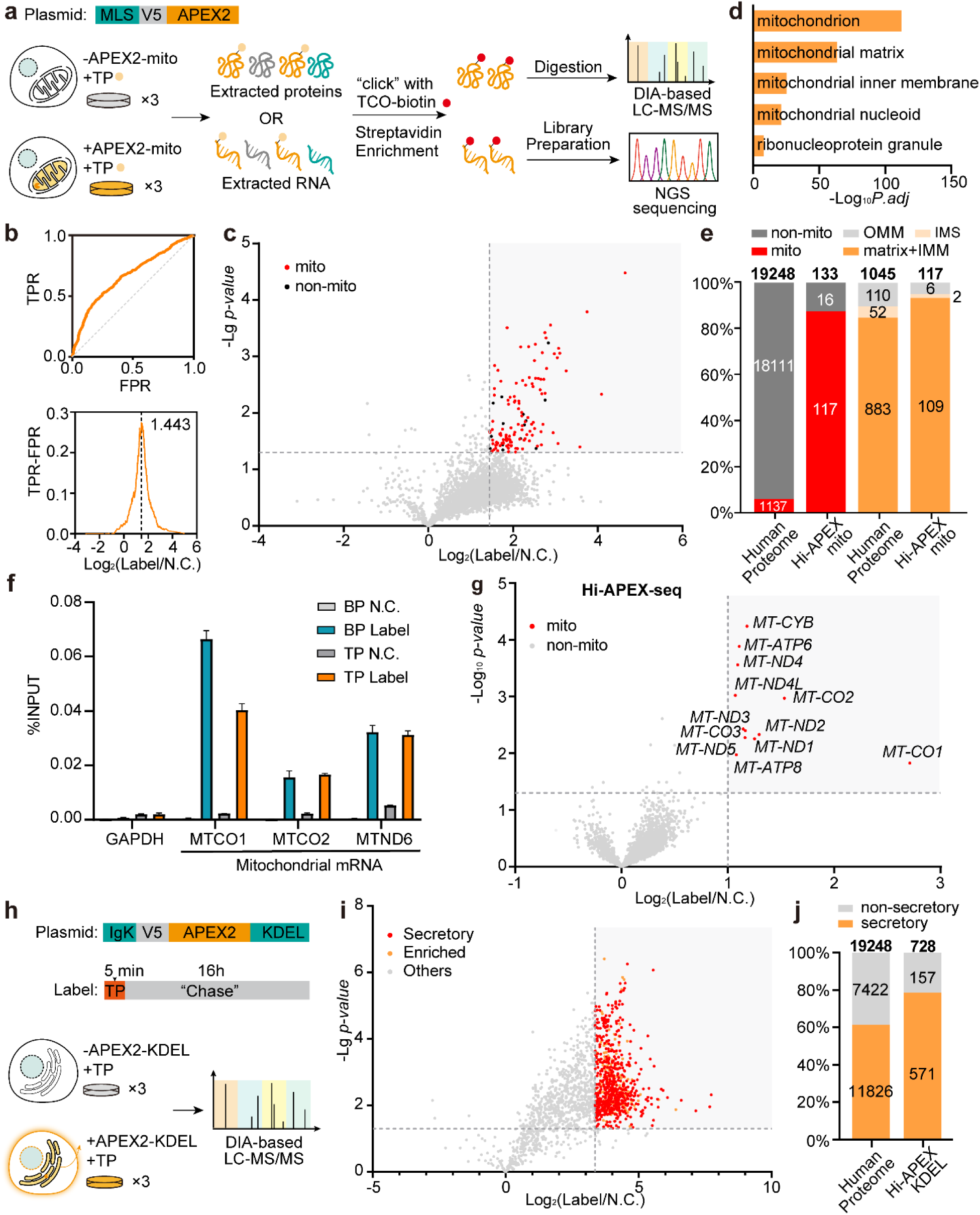
Benchmarking Hi-APEX for spatial proteomics and transcriptomics. (a) Schematic of the Hi-APEX strategy for spatial proteomic and transcriptomic mapping of the mitochondrial matrix, including the APEX2-mito plasmid design. (b) ROC analysis of protein enrichment in Hi-APEX-mito proteomics versus an unlabeled control. True positives are defined as known mitochondrial proteins; false positives are non-mitochondrial proteins. (c) Volcano plots depicting protein enrichment from Hi-APEX-mito labeling compared to an unlabeled control. (d) Gene Ontology Cellular Component (GOCC) analysis of proteins enriched by Hi-APEX-mito. (e) Specificity analysis of Hi-APEX-mito-enriched proteins for mitochondrial and sub-mitochondrial localization. (f) Enrichment of mtDNA-encoded transcripts detected by quantitative RT-PCR. H₂O₂-dependent BP labeling served as a positive control. (g) Volcano plots showing transcript enrichment from mitochondrial matrix-localized Hi-APEX-seq compared to an unlabeled control. (h) Schematic of Hi-APEX pulse-chase labeling for dynamic mapping of protein secretion from the ER, illustrating the APEX2-KDEL plasmid and experimental workflow. (i) Volcano plots showing enrichment of secreted proteins identified by Hi-APEX-KDEL over an unlabeled control. (j) Enrichment levels of known secreted proteins captured by the Hi-APEX-KDEL method.

Existing spatial transcriptomics methods (APEX-seq^10^, photocatalytic labeling^18,28^) induce cellular stress and are incompatible with living organisms due to H₂O₂ toxicity or limited tissue penetration of visible light. We hypothesized that TP-generated radicals could also label RNA, enabling Hi-APEX-seq as a non-toxic, *in vivo*-compatible spatial transcriptomics method. Using the same APEX2-mito samples, we extracted RNA, performed IEDDA conjugation with TCO-biotin, and conducted streptavidin pulldown (**Fig. 3a**). Streptavidin dot blot confirmed APEX-dependent RNA labeling (**Supplementary Fig. 3b**). qRT-PCR revealed significant enrichment of mitochondrial transcripts (MT-CO2, MT-ND6) over cytosolic mRNAs (**Fig. 3f**). Transcriptome-wide sequencing (Hi-APEX-seq) showed high reproducibility (**Supplementary Fig. 3c**) and specifically enriched mtDNA-encoded mRNAs and rRNAs, while nuclear-encoded RNAs remained unenriched (**Fig. 3g, Supplementary Fig. 3d, Table S2**). These results demonstrate Hi-APEX’s capability for simultaneous spatial proteomics and transcriptomics.

We next applied Hi-APEX to map proteome trafficking using a pulse-chase design—an application incompatible with traditional APEX (**Fig. 3h**). After ER luminal labeling (APEX2-KDEL) and TP removal, cells were cultured for 16 hours before secretome collection. We observed labeling of secreted proteins that trafficked from the ER during the chase period (**Supplementary Fig. 4a**). DIA analysis of labeled secretomes identified 728 ER-originating secreted proteins (**Fig. 3i, Supplementary Fig. 4b, Table S3**), significantly enriching extracellular and known secretory proteins (**Fig. 3j, Supplementary Fig. 4c**). Taken together, Hi-APEX demonstrates high specificity in resolving spatiotemporal distribution of biomolecules.

## Simultaneous proteomic and transcriptomic mapping of stress granules using Hi-APEX

Stress granules (SGs) are dynamic, membraneless organelles that form in response to cellular stress. Since their composition is highly stress-dependent, mapping their authentic molecular landscape requires methods that introduce minimal perturbation. Conventional APEX2-based profiling relies on H₂O₂, an exogenous oxidative stressor that may itself alter SG content^29,30^. To address this, we developed Hi-APEX for the simultaneous mapping of the SG proteome and transcriptome under native stress conditions (**Fig. 4a**). We fused APEX2 to the core SG protein G3BP1 in HEK293T cells. SG formation was induced with sodium arsenite, followed by brief Hi-APEX labeling. This resulted in specific labeling of SGs, visualized by co-localization with the SG marker G3BP2, while labeling was diffuse in the cytosol under basal conditions (**Fig. 4b**). Critically, the Hi-APEX workflow did not enhance SG formation, confirming minimal perturbation (**Supplementary Fig. 5a**). Streptavidin blotting confirmed the enzyme-dependent labeling with Hi-APEX-G3BP1(**Supplementary Fig. 5b**).

**Fig. 4.**
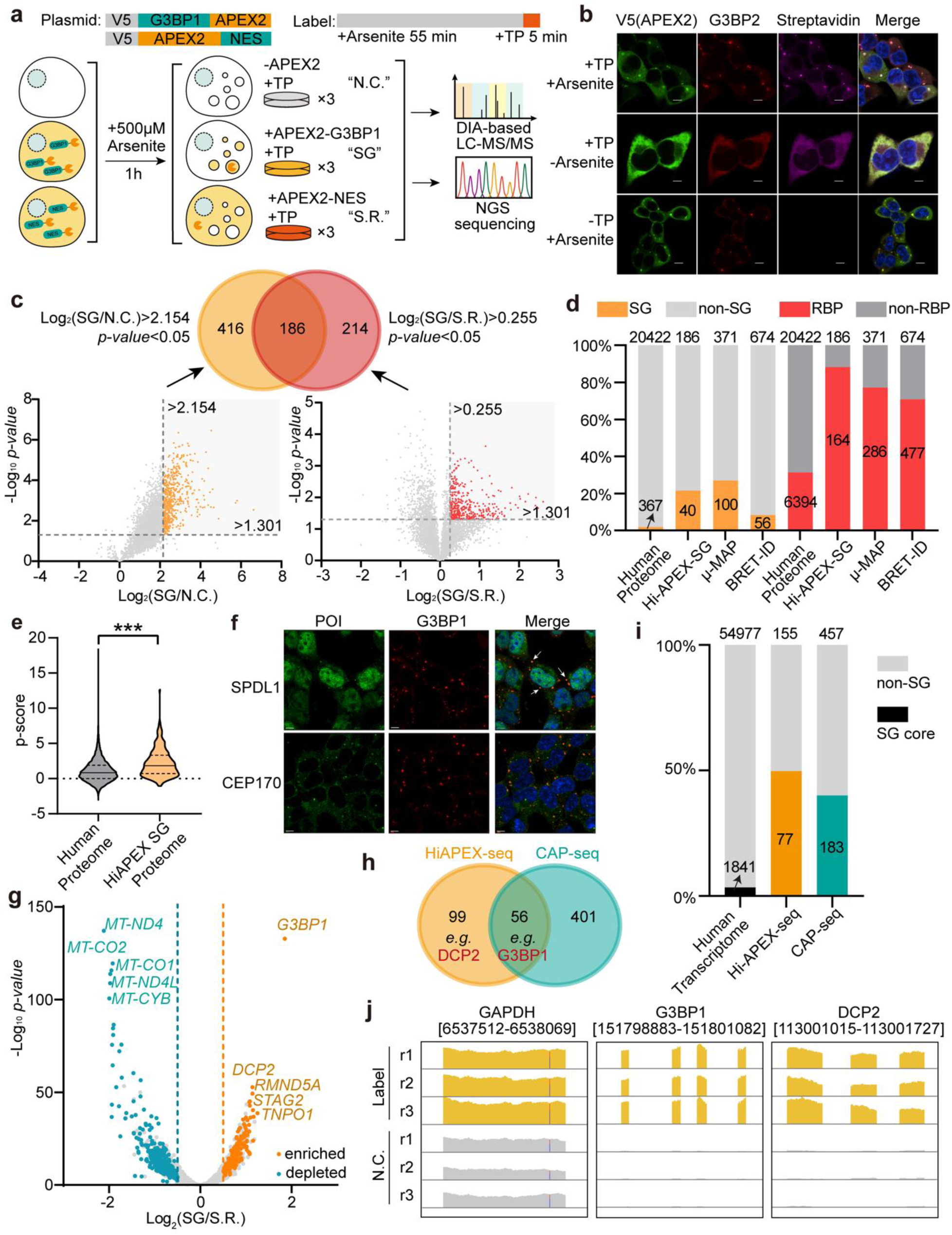
Simultaneous proteomic and transcriptomic mapping of stress granules using Hi-APEX. (a) Schematic of Hi-APEX for spatial proteomic and transcriptomic mapping of stress granules (SGs), showing the APEX2-G3BP1 (SG-targeted) and APEX2-NES (cytosolic control) plasmid constructs. Cells lacking APEX2 expression served as the negative control (N.C.), while those expressing cytosolic APEX2-NES provided the spatial reference (S.R.). (b) Confocal microscopy of Hi-APEX labeling in SGs. APEX2 was detected with an anti-V5 antibody, SGs were marked with an anti-G3BP2 antibody, and biotin incorporation from TP labeling was visualized with streptavidin. Scale bars, 10 µm. (c) Protein enrichment criteria for the SG proteome. Labeled samples from SG-targeted Hi-APEX were compared to both the N.C. and S.R. Proteins surpassing ROC-defined enrichment thresholds with a *p*-value < 0.05 in both comparisons were classified as SG-enriched. (d) Enrichment of known SG and RNA-binding proteins identified by SG-targeted Hi-APEX, compared to the established methods µMAP and BRET-ID. (e) Phase separation propensity (P-scores) of proteins enriched by SG-targeted Hi-APEX. \*\*\**p* < 0.001. (f) Confocal images showing the colocalization of SPDL1 and CEP170 with SGs following arsenite treatment. Scale bars, 10 µm. (g) Volcano plot showing transcript enrichment from SG-localized Hi-APEX-seq over the cytosolic spatial reference (S.R.). (h) Overlap of transcripts enriched by SG-localized Hi-APEX-seq with those identified by CAP-seq. (i) Proportion of known SG core transcripts captured by SG-localized Hi-APEX-seq. (j) Integrative genomics viewer (IGV) plots showcasing the enrichment of DCP2, G3BP1 and GAPDH transcripts by Hi-APEX-seq.

Proteomic analysis of Hi-APEX-G3BP1 labeling, compared against non-labeled and cytosolic controls, revealed significant enrichment of known SG core components, including G3BP1, CAPRIN1, PRRC2C, and UBAP2 (**Fig. 4c**). Following ROC and statistical filtering, we identified 186 high-confidence SG proteins and their GO analysis reveals significant enrichment of stress granule assembly (**Supplementary Fig. 5c-e and Table S4**). This set was highly enriched for established SG and RNA-binding proteins, performing comparably to (or better than) the high-resolution method µMAP^31^ and BRET-ID^32^ (**Fig. 4d**), and exhibited significantly higher phase separation propensity (**Fig. 4e**). We validated two novel “SG orphans”, Protein Spindly (SPDL1) and Centrosomal protein of 170 kDa (CEP170), confirming their recruitment to arsenite-induced SGs (**Fig. 4f**).

In parallel, we performed Hi-APEX-seq on the same samples to define the SG transcriptome, identifying 155 enriched RNAs (**Supplementary Fig. 5f-g and Table S5**). The G3BP1 mRNA itself was the most significantly enriched transcript (**Fig. 4g**). This Hi-APEX-seq dataset complements the CAP-seq^33^ results, as the latter employs additional stressors (e.g., blue light) that likely recruit a wider array of RNAs (**Fig. 4h**). Indeed, Hi-APEX-seq exhibits higher enrichment of SG core transcripts^34^ (**Fig. 4i**), uniquely capturing key clients like DCP2 mRNA, whose protein product is associated with P-bodies^35^ (**Fig. 4j**). GO analysis of these transcripts exhibits enrichment of biological processes including regulation of microtubule cytoskeleton organization and protein ubiquitination (**Supplementary Fig. 5h**). Together, Hi-APEX provides a minimally perturbative PL method for spatial multi-omic profiling, enabling the identification of the authentic molecular composition of subcellular compartments.

## Mapping the mitochondrial GPx4 interactome during ferroptosis via Hi-APEX

Glutathione peroxidase 4 (GPx4) is the central regulator of ferroptosis, a non-apoptotic cell death driven by lipid peroxidation^36^. While the cytosolic GPx4 isoform is a well-established ferroptosis inhibitor, the role of the mitochondrial isoform (mGPx4) remains controversial^37,38^. We hypothesized that mapping the protein interactors of mGPx4 could reveal its functional networks and regulatory mechanisms during ferroptosis—a process involving dynamic mitochondrial changes^39^. We therefore applied Hi-APEX to define the mGPx4 interactome under ferroptotic conditions (**Fig. 5a**), as conventional APEX2 labeling is precluded for studying ferroptosis due to the requirement for H₂O₂, a key driver of this cell death pathway^40^.

**Fig. 5.**
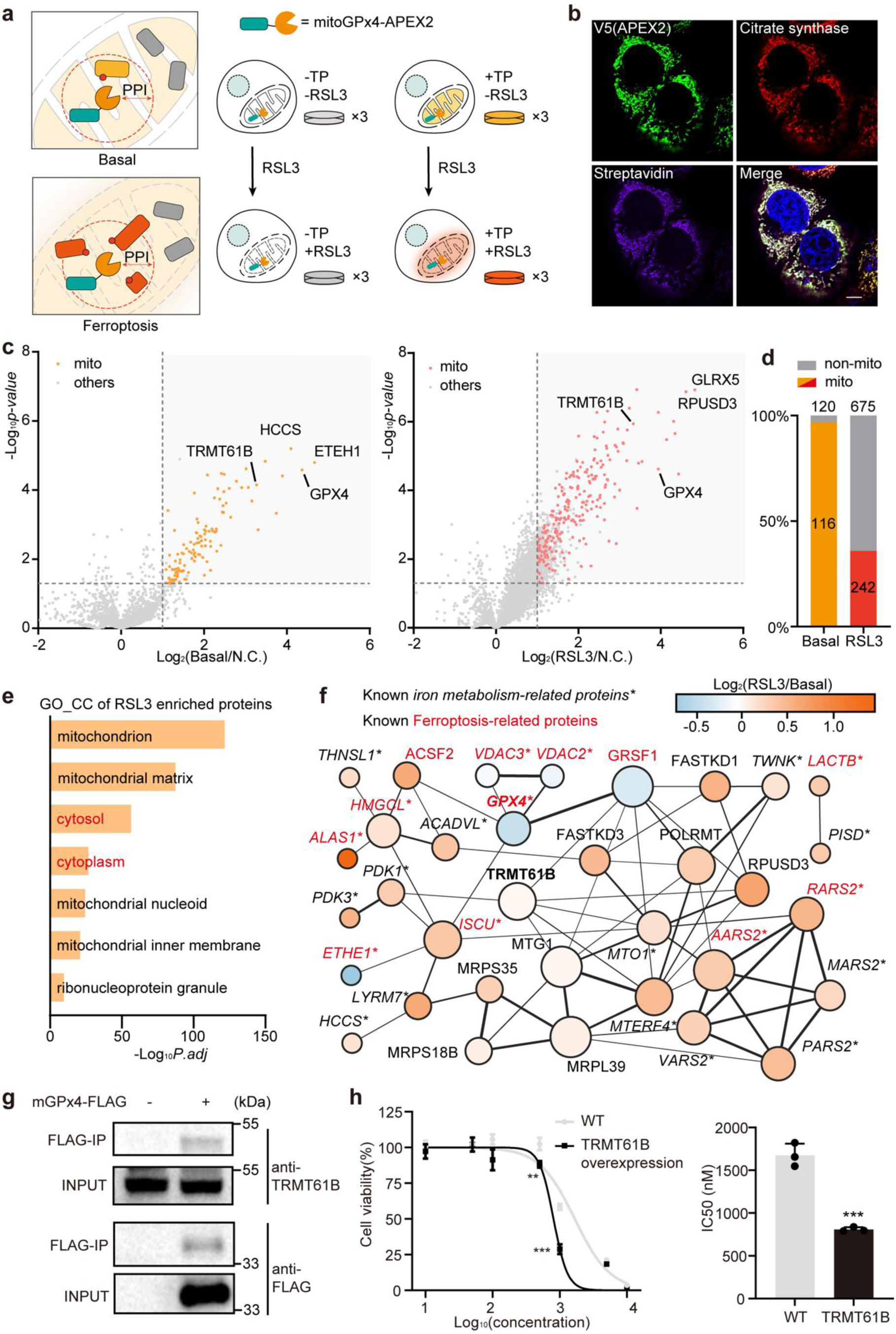
Mapping the mitochondrial GPx4 interactome during ferroptosis via Hi-APEX. (a) Schematic of Hi-APEX for interactome mapping of mitochondria-localized GPx4 (mGPx4) under basal and RSL3-induced ferroptosis conditions. (b) Confocal microscopy of Hi-APEX-mGPx4 labeling. APEX2 was detected with an anti-V5 antibody, mitochondria were labeled with an anti-citrate synthase antibody, and biotin incorporation from TP labeling was visualized with streptavidin. Scale bars, 10 µm. (c) Volcano plots showing protein enrichment by Hi-APEX-mGPx4 over an unlabeled control under basal (left) and RSL3-treated (right) conditions. (d) Mitochondrial specificity of proteins enriched by Hi-APEX-mGPx4 under basal (left) and RSL3-treated (right) conditions. (e) GOCC analysis of proteins enriched by Hi-APEX-mGPx4 following RSL3 treatment. (f) STRING network analysis of the top-enriched mGPx4 interactors identified by Hi-APEX. (g) Validation of TRMT61B as an mGPx4-interacting protein by co-immunoprecipitation. Lysates from HT1080 cells stably expressing Flag-tagged mGPx4 were subjected to anti-Flag immunoprecipitation and probed for TRMT61B. (h) Effect of TRMT61B overexpression on RSL3-induced ferroptosis. Cell viability was assessed by MTS assay in wild-type (WT) and TRMT61B-overexpressing HT1080 cells after a 3-hour treatment with increasing RSL3 concentrations. Calculated IC₅₀ values are shown. \*\**p* < 0.01; \*\*\**p* < 0.001; n.s., not significant. Data represented as mean ± SD.

We generated HT1080 cells stably expressing APEX2-tagged mGPx4, confirming mitochondria-specific TP labeling (**Fig. 5b; Supplementary Fig. 6a**). Ferroptosis was induced with RSL3 without interference from the Hi-APEX workflow (**Supplementary Fig. 6b**). Proteomic analysis identified 120 and 675 significantly enriched proteins in basal and ferroptotic conditions, respectively, with GPx4 as the top hit (**Fig. 5c; Supplementary Fig. 6c; Table S6**). Over 97% of basal interactors were mitochondrial, confirming high spatial specificity (**Fig. 5d**). In contrast, the ferroptotic interactome included substantial cytosolic proteins enriched for phosphorylation and ubiquitination pathways, suggesting their recruitment to mitochondria, potentially due to increased membrane permeability (**Fig. 5e; Supplementary Fig. 6d**).

The top-enriched interactors were predominantly iron and redox metabolism proteins, forming extensive networks (**Fig. 5f; Supplementary Fig. 6e**). Key findings included decreased interaction with the ferroptosis suppressor ETHE1^41^, and increased association with known regulators like GLRX5^42^, ALAS1^43^, and ACSF2^44^, indicating mGPx4 coordinates ferroptosis through a broad network.

The prevalence of known ferroptosis regulators among our high-confidence hits led us to hypothesize that other identified interactors might also influence this process. GO analysis indicated a significant enrichment for proteins involved in mitochondrial transcription and translation (**Supplementary Fig. 6f**), including the m¹A modifier TRMT61B that mediates m¹A modifications on mitochondrial tRNAs to regulate translational efficiency^45^. To biochemically validate their interactions, we performed co-immunoprecipitation with Flag-tagged mGPx4 in HT1080 cells, which confirmed the association of TRMT61B with mGPx4 (**Fig. 5g**). To assess their functional role in ferroptosis, we treated cells overexpressing TRMT61B with titrating doses of RSL3 (**Supplementary Fig. 6g**). This functional assay revealed that elevated TRMT61B expression sensitized the cells to ferroptosis (**Fig. 5h**), suggesting potential links between mitochondrial RNA epigenetics and the regulation of cell death.

## *In vivo* Hi-APEX labeling in tumor xenografts and hippocampal neurons

Following the successful validation of Hi-APEX in cell culture, we exploited its intrinsic *in vivo* compatibility to label and identify mGPx4-proximal proteins in tumor xenografts. We subcutaneously injected nude mice with HT1080 cells stably expressing mGPx4-APEX2, allowing tumors to form over two weeks before performing intratumoral TP injection for *in vivo* labeling (**Fig. 6a**). Streptavidin blotting of tumor lysates after TCO-biotin conjugation revealed robust, APEX2-dependent protein labeling, with no signal detected in wild-type tumors (**Fig. 6b**). We then isolated the Hi-APEX-labeled proteins from tumor tissue lysates using streptavidin beads for proteomic analysis, which identified a total of 228 high-confidence mGPx4-proximal proteins (**Fig. 6c; Supplementary Fig. 7a and Table S7**). These included known interactors from cell culture, such as ETHE1, as well as proteins unique to the tumor xenograft environment, like the reactive oxygen species modulator ROMO1^46^. Consistent with our cell culture profiling, GOCC analysis revealed significant enrichment of mitochondrial proteins, and “mitochondrial translation” emerged as the most enriched biological process (**Fig. 6d**). These findings demonstrate that Hi-APEX is fully compatible with *in vivo* applications and can uncover interaction heterogeneities between cultured cells and native tissue environments.

**Fig. 6.**
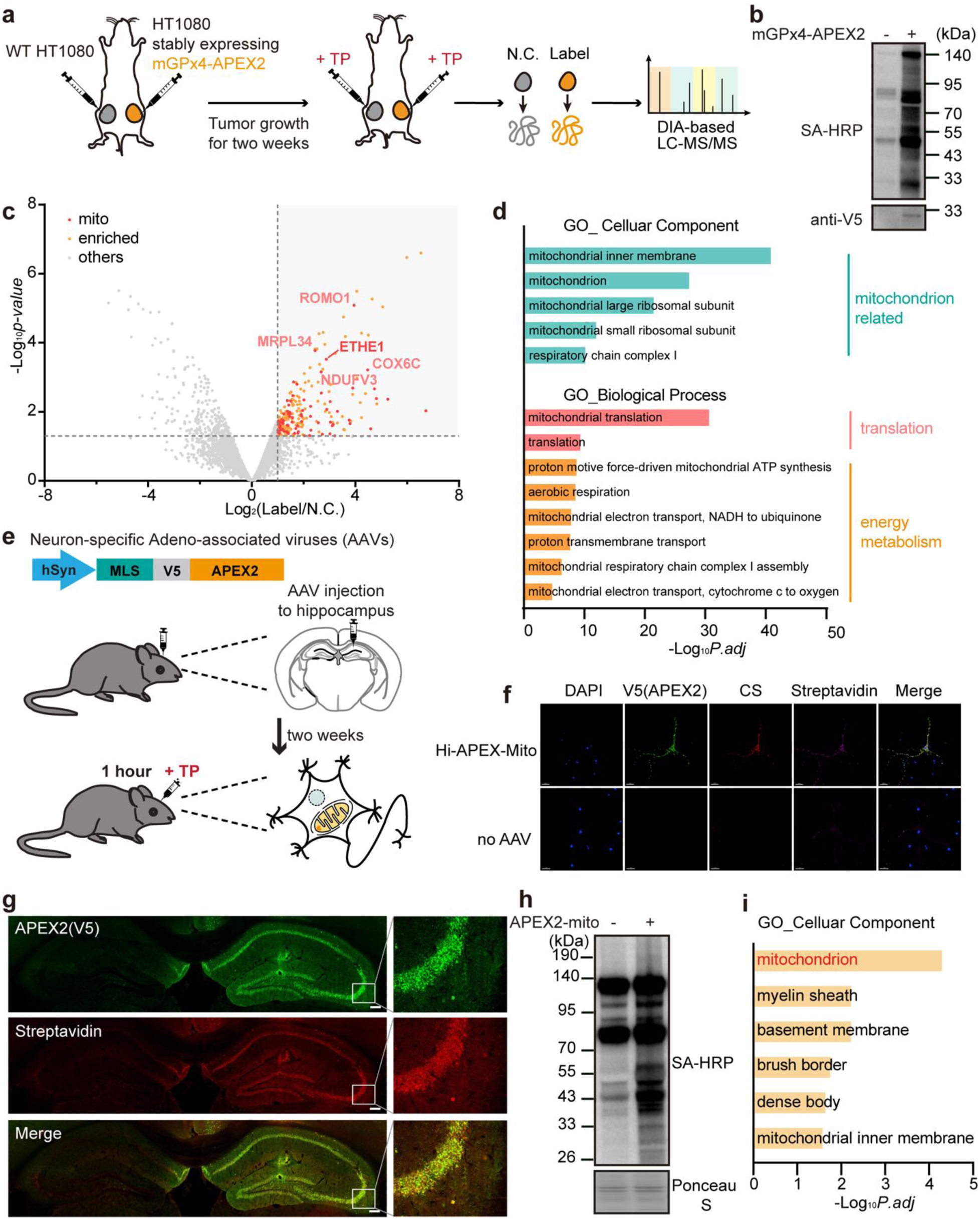
*In vivo* Hi-APEX labeling in tumor xenografts and hippocampal neurons. (a) Schematic of *in vivo* Hi-APEX labeling. Mice bearing HT1080 tumors expressing mGPx4-APEX2 received intratumoral injections of TP. (b) Streptavidin blot analysis of mGPx4-targeted Hi-APEX labeling in tumor xenografts. Labeled tumor tissues were lysed, conjugated with TCO-biotin via IEDDA, and analyzed by streptavidin blotting. (c) Volcano plots showing enrichment of proteins identified by Hi-APEX-mGPx4 over an unlabeled control *in vivo*. (d) GOCC analysis of mGPx4-interacting proteins identified by *in vivo* Hi-APEX. (e) Schematic of *in vivo* Hi-APEX labeling in mouse hippocampal neurons. Adeno-associated viruses (AAVs) encoding APEX2 targeted to the mitochondrial matrix were injected into the hippocampus. After two weeks for expression, TP was injected into the same site for 1-hour labeling. (f) Confocal microscopy of Hi-APEX-mito labeling in cultured mouse cortical neurons. Neurons were infected with AAVs expressing APEX2-mito for 10 days, then labeled with 500 µM TP for 5 minutes. APEX2 was visualized with an anti-V5 antibody, mitochondria with anti-citrate synthase, and biotin incorporation with streptavidin. Scale bars, 10 µm. (g) Immunofluorescence staining of hippocampal sections with or without TP treatment. APEX2 was detected with an anti-V5 antibody, and biotin labeling was visualized with streptavidin. Scale bars, 50 µm. (h) Streptavidin blot analysis of mitochondrial matrix-targeted Hi-APEX labeling from the hippocampi of living mice. The labeled hippocampal regions were dissected, lysed, conjugated with TCO-biotin, and analyzed. (i) GOCC analysis of proteins enriched by *in vivo* Hi-APEX labeling from the mitochondrial matrix of hippocampal neurons.

Finally, we applied Hi-APEX to map the mitochondrial matrix proteome within hippocampal neurons of the intact mouse brain (**Fig. 6e**). While peroxidase-based PL methods, such as APEX2 and HRP, have been used in the central nervous system, these studies were conducted *ex vivo* due to the toxicity of direct H₂O₂ injection ^47–49^. Hi-APEX overcomes this critical limitation, enabling genuine *in vivo* APEX2 labeling. We packaged a mitochondria-targeted APEX2 construct, under a neuronal promoter, into adeno-associated virus (AAV) particles. We first confirmed functional AAV expression and Hi-APEX labeling in mouse cortical neurons, where streptavidin blotting showed robust protein biotinylation after a 1-hour TP treatment (**Supplementary Fig. 7b**). Confocal imaging confirmed precise colocalization of the TP labeling signal with both APEX2 and mitochondrial markers, validating the high spatial specificity of Hi-APEX in cultured neurons (**Fig. 6f**).

We then injected an AAV encoding APEX2-mito into the mouse hippocampus and allowed two weeks for enzyme expression. For initial Hi-APEX validation, we extracted the brains, sectioned them into 300 µm slices in artificial cerebrospinal fluid (ACSF), and incubated the slices with 500 µM TP for 30 minutes (**Supplementary Fig. 7c**). Streptavidin blotting confirmed enzyme-dependent TP labeling, and DIA-based proteomics demonstrated high mitochondrial specificity among the enriched proteins (**Supplementary Fig. 7d,e**). To achieve true *in vivo* labeling, we instead administered TP directly at the same injection site and allowed the labeling to proceed for 1 hour in the live animal (**Fig. 6e**). On subsequent brain sections, an IEDDA reaction with TCO-biotin followed by streptavidin-Alexa Fluor 647 staining revealed robust labeling that was highly enriched in the hippocampus and colocalized with APEX2 expression (**Fig. 6g**). Control experiments (omitting either AAV or TP) showed only minimal background. Furthermore, streptavidin blotting of hippocampal lysates confirmed extensive protein labeling specifically in TP-treated samples, with negligible signal in controls (**Fig. 6h**). Subsequent DIA-based proteomics of the streptavidin-enriched fractions identified the labeled proteins, which were significantly enriched for mitochondrial components (**Fig. 6i and Table S8**). Collectively, these results establish that Hi-APEX enables straightforward *in vivo* PL through simple TP injection.

## Discussion

PL techniques, such as APEX2, have revolutionized cellular proteomics by enabling the spatial mapping of protein interactions^5^. Conventional APEX2 functions by using exogenous H₂O₂ to catalyze biotin-phenol into reactive phenoxyl radicals that tag nearby proteins^6^. While powerful, this requirement for high concentrations of H₂O₂ induces significant cellular toxicity and stress, which can confound experimental results, especially in sensitive models. We address this fundamental limitation with the discovery of a tetrazine-phenol probe that APEX2 activates through an H₂O₂-independent mechanism. This new method, which we call Hi-APEX, achieves labeling specificity and efficiency comparable to the conventional system across multiple subcellular regions, but without the associated oxidative damage.

The elimination of exogenous H₂O₂ allows Hi-APEX to be applied in contexts where minimal perturbation is critical. We successfully demonstrated its utility in three such challenging scenarios: mapping protein secretion dynamics via pulse-chase labeling, conducting spatial multi-omics on stress granules under authentic oxidative stress, and profiling the interactome of mitochondria-localized GPX4 during the dynamic process of ferroptosis^36,37,40^. These applications generated rich datasets that led to the validation of novel biology, including the confirmation of new stress granule residents and a functional interaction between mGPX4 and TRMT61B. This interaction was found to promote ferroptosis, likely through TRMT61B’s role in regulating m¹A modifications on mitochondrial tRNA.

A paramount advantage of Hi-APEX is its capacity for genuine *in vivo* PL. Existing APEX2 models, such as transgenic mice or AAV-delivered constructs, still require tissue dissection and *ex vivo* H₂O₂ treatment to initiate labeling. Hi-APEX overcomes this barrier, enabling direct labeling within living mice, as we demonstrated by mapping the mGPX4 interactome in tumor xenografts and quantifying the mitochondrial matrix proteome in hippocampal neurons.

Furthermore, Hi-APEX avoids the key drawbacks of other PL methods for *in vivo* applications, such as high background (TurboID)^14^, poor tissue penetration (photocatalytic methods)^20^, or copper toxicity (TyroID^15,16^/LaccID^17^). Hi-APEX-seq in particular surpasses existing techniques like APEX-seq^10,50^, CAP-seq^18^, and µMap-seq^28^, positioning itself as the only PL method currently available for *in vivo* spatial transcriptomics. Crucially, it requires no enzyme engineering, allowing for immediate adoption in any existing APEX2 system.

While the precise mechanism of H₂O₂-independent radical generation by the TP probe requires further elucidation, our initial evidence suggests the formation of an H₂O₂-like intermediate. A full understanding will necessitate further computational and experimental validation.

Determining the co-crystal structure of APEX2 bound to TP will be particularly important for clarifying the catalytic role of residues like His42 and for informing the design of more potent next-generation probes.

Finally, Hi-APEX offers a transformative path for extending dynamic proteomic technologies like our previously developed TransitID^22^. TransitID, which tracks protein trafficking by combining TurboID and APEX2, has been confined to cell culture due to the toxicity of conventional APEX2’s *in vivo* labeling step. Because Hi-APEX uses a non-biotin probe that is orthogonal to TurboID, it serves as a perfect replacement. This compatibility opens the door to an *in vivo* TransitID platform, capable of elucidating proteome trafficking between non-dissectible cell types within intact living systems and unlocking novel biological insights currently inaccessible to other technologies.

## Acknowledgements

We would like to thank the assistance of Imaging Core Facility of the Protein Research Center for Technology Development, Tsinghua University. This work was supported by the National Natural Science Foundation of China (92478128 and 22477066 to W.Q., 22407141 to Y.C.), the National Key Research and Development Program of China (No. 2024YFA1308000, W.Q.), the Chinese Academy of Medical Sciences (CAMS) Innovation Fund for Medical Sciences (2024-I2M-3-010, Y.C.), Beijing Natural Science Foundation (JQ25018, W.Q.), Youth Talent Cultivation Fund of Tsinghua University (W.Q.), “Dushi Plan” from Tsinghua University (W.Q.), Beijing Frontier Research Center for Biological Structure, the Fundamental Research Funds from Beijing National Laboratory for Molecular Sciences (BNLMS202301, W.Q.) and the Shenzhen Medical Research Fund (B2401004, W.Q.). W.Q. is supported by Bayer Investigator Award.

## Author contributions

B.C., H.G., Y.C., and W.Q. designed the research and analyzed all the data except where noted. B.C. and H.G. performed all experiments except where noted. Z.Y. evaluated Hi-APEX *in vitro* and in cells. W.L. performed Hi-APEX labeling in tumor tissues. C.L. performed Hi-APEX labeling in mouse brains under the supervision of Z.W. S.X. solved the crystal structure of APEX2. Y.Z. performed cell viability assays. H.G. performed Hi-APEX labeling in cultured neurons. S.S. performed mGPx4 co-IP experiments. X.S. performed imaging validation of SG proteins. S.Z. performed molecular docking under the supervision of L.L. SG.Q. analyzed RNA sequencing data under the supervision of Y.C. B.C., H.G., Y.C., and W.Q. wrote the paper with inputs from all authors.

## Declaration of interests

The authors declare no competing interests.

## Online methods

### Cell culture

HEK293T, HeLa and HT1080 cells from the ATCC (passages <25) were cultured in DMEM (Thermo Fisher) supplemented with 10% fetal bovine serum (Biological Industries), 100 units/mL penicillin, and 100 mg/mL streptomycin at 37 °C under 5% CO_2_. For fluorescence microscopy imaging experiments, cells were grown on 15-mm glass-bottom cell culture dish (NEST). For proteomic and transcriptomic experiments, cells were grown on 10-cm glass-bottomed Petri dishes (NEST). For Western Blot experiments, cells were grown on six-well plates (NEST).

For primary cortical neuron culture, primary cortical neurons were isolated from postnatal day 0 (P0) mouse pups. Following sterilization with 75% ethanol and decapitation, the brains were extracted and placed in ice-cold serum-free medium. Under a stereomicroscope, the cortical hemispheres were dissected, and the meninges, blood vessels, hippocampi, and subcortical tissues were meticulously removed. The cleaned cortical tissues were then rinsed twice with ice-cold PBS and digested in pre-warmed 0.25% Trypsin-EDTA at 37°C for 15-20 minutes. The digestion was terminated by adding an equal volume of complete DMEM, and the tissue was mechanically dissociated into a single-cell suspension via gentle trituration. The suspension was passed through a 70 µm cell strainer and centrifuged at 500 x g for 5 minutes. After discarding the supernatant, the cell pellet was resuspended in serum-free medium, and the cells were counted using a hemocytometer. Neurons were seeded into 6-well plates at a density of 6-8 × 10⁵ cells per well. After 3-6 hours of incubation to allow for cell attachment, the cultures were gently washed once with pre-warmed PBS, and the medium was completely replaced with pre-warmed NBPG medium (Neurobasal medium supplemented with 2% B-27 Supplement, 1% GlutaMAX, and 1% Penicillin-Streptomycin). Subsequently, the culture medium was fully replaced with fresh, pre-warmed NBPG medium every 1.5 days.

### Plasmids

Polymerase chain reactions (PCR) were performed with Phanta DNA Polymerase (catalog no. P520) purchased from Vazyme. Ligase-free cloning reactions were performed with Basic Seamless Cloning and Assembly Kit (catalog no. CU201) purchased from TransGen. Plasmid products were transformed into DH5α competent cell (catalog no. TSC-C14) purchased from Tsingke. DNA sequences were confirmed by Sanger sequencing before use.

pLX304-Mito-V5-APEX2 was a gift from Dr. Chu Wang at Peking University. The other plasmids (APEX2-NES, APEX2-NLS, APEX2-OMM, APEX2-ERM, APEX2-KDEL, APEX2-G3BP1, mGPx4-APEX2) of APEX2 were constructed from pLX304-Mito-V5-APEX2 via PCR. The plasmids contain the DNA sequence of mGPx4-FLAG and TRMT61B were ordered from MIAOLING Biology (catalog no. P46156, P3553, MIAOLING Biology)

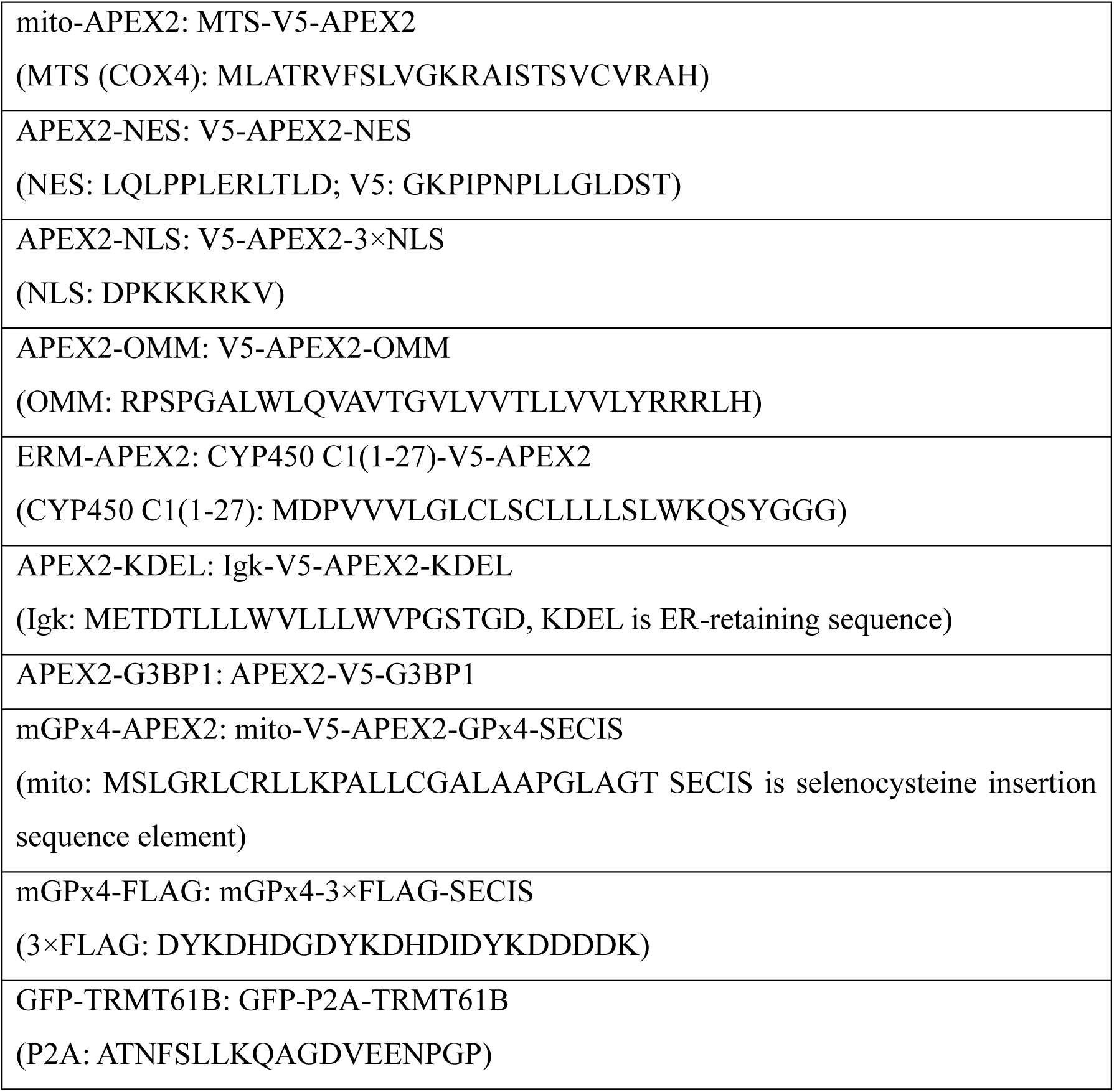

APEX2-TM was constructed from HRP-TM, which was purchased from Addgene (Plasmid #44441) by replacing HRP with APEX2.

### Protein purification

APEX2 gene was cloned into a modified pQLinkHx vector with a N-terminal 8× His tag, and the plasmid was introduced into E. coli BL21(DE3) for protein expression. The E. coli cells were cultured in 2 × YT at 37°C with 200 r.p.m. until the OD_600_ reached 0.6, and induced by adding isopropyl-β-D-thiogalactopyranoside (IPTG) to a final concentration of 0.2 mM and final concentration 1mM 5-ALA continued incubating at 16°C for overnight.

For APEX2 protein purification, cells were re-suspended in lysis buffer (50 mM Tris–HCl pH 8.0, 200 mM NaCl, 2.5% (v/v) glycerol, 2 mM PMSF, protease inhibitor cocktail (Roche)) and lysed using a high-pressure cell disrupter at 900 bar for 5 min. Cell debris was removed by centrifugation at 17,418 × g for 1 h, and the supernatant was supplemented with Hemin (catalog no.H886057, MACKLIN) to a final concentration of 0.32 mM, followed by incubation at 4°C overnight. The supernatant after incubation was loaded to a Ni-NTA gravity column, Unbound proteins were washed away with lysis buffer containing 50 mM imidazole, and APEX2 protein was eluted with lysis buffer containing 250 mM imidazole. Then, the 8 × His tag was cleaved by using Tev treatment at 4 ℃ for overnight. The digested product was then loaded to a pre-equilibrated 5-ml HisTrap Ni-NTA column (Cytiva), and flow-through was purified by gel filtration (Superose 75 Increase 10/300 GL, GE Healthcare) pre-equilibrated in buffer B (20 mM Tris–HCl pH 7.5, 150 mM NaCl,) supplemented with 1 mM TCEP. The elution fractions were analysed by SDS–PAGE, and the peak fractions of the purified APEX2 protein were collected and concentrated for *in vitro* labeling and crystallization.

### Crystallization and structure determination

Purified APEX2 was concentrated to 12 mg/mL. Crystals were obtained by mixing the proteins with equal volume reservoir buffer containing 1.6 M MgSO_4_, 0.1 M MES, pH 6.5. The proteins were crystallized by sitting drop vapor diffusion at 18℃. The crystals were flash-frozen in liquid nitrogen using nylon loops after removing excess mother liquor. Diffraction data for the APEX2 were collected on a Rigaku XtaLAB SynergyCustom detector at the X-ray Crystallography Platform, School of Life Sciences, Tsinghua University. The structures were solved by the molecular replacement method using PHASER^51^ with PDB entry 1OAF^25^ as the search model. Crystallographic refinements were performed using PHENIX version 1.14^52^. Manual rebuilding and adjustment of the structures were done in Coot^53^.

### Molecular docking

Molecular docking studies were conducted using the Glide^54^ module of the Schrödinger Suite (release 2022-1). The protein structure was derived from X-ray crystallography and prepared using the Protein Preparation Wizard, which included the addition of missing hydrogen atoms, optimization of hydrogen bonding networks via PROPKA^55^ at pH 7.0, and restrained energy minimization with the OPLS4 force field^56^ using an RMSD convergence threshold of 0.3 Å. The TP and BP ligands from 2D to 3D were geometry-optimized in the LigPrep module using the OPLS4 force field. Protonation and ionization states, as well as stereoisomers, were enumerated using Epik^57^ at pH 7.0 ± 2.0. A receptor grid was generated using the Glide Receptor Grid Generation tool. An inner box (ligand diameter midpoint box) of 15 × 15 × 15 Å was centered around the HEM binding site. Docking was performed using the Extra Precision (XP) protocol, with flexible ligand sampling enabled to improve the accuracy of binding pose predictions. The docking pose with the highest score was subsequently analyzed.

### Surface plasmon resonance

Surface plasmon resonance (SPR) experiments were conducted on a Biacore 8k + system using CM5 sensor chips (GE Healthcare). The CM5 surfaces were activated at 25°C by injecting a mixture of 0.4 M EDC and 0.1 M NHS (Cytiva) at a flow rate of 10 μl min^-1^ for 420 s. The APEX2 protein purified under PBS were then diluted to a concentration of 200 μg mL^-1^ in sodium acetate (pH5.0), and injected separately at a flow rate of 10 μL min^-1^ until a density of approximately 15,000RU of immobilized APEX2 was achieved, To block any remaining activated sites, a 420-s injection of ethanolamine (1 M, pH 8.5)at a flow rate of 10 μL min^-1^ was performed.

To measure the binding activity of small molecules to proteins, Serial dilution gradients (6.25-1600 nM) of the BP, AP, TP were flowed through with a running buffer containing 10 mi HEPES, 150 mM Nacl, pH 7.2, 0.05% (v/v Tween 20., The association time was set to 90 s, followed by a dissociation time of 90 s. The obtained data were analyzed using Biacore Insight Evaluation Software (GE Healthcare). to determine the affinity.

### Construction of stable cell lines

The constructed pLx304 plasmids were co-transfected with pMD2.G and pSPAX2 into HEK293T cells using neofect Transfection Reagent (catalog no. TF201201, Neofect). Viral supernatant was collected 48 hours after transfection and transduced to the indicated HEK293T or HT1080 cell line for 48 hours. Cells were selected using the complete culture medium with 5 μg/mL Blasticidin S (catalog no. HY-103401, MCE) for approximately 1 week, subsequently maintained in the complete culture medium.

### Hi-APEX labeling *in vitro* and in living cells

For the labeling of Hi-APEX on BSA, 2 mg/mL BSA was incubated with 2 μM APEX2 or HRP and 500 μM TP at 37℃ for 30 min. The reaction was stopped by precipitating the BSA. The precipitated BSA was resuspended in 0.4% SDS/PBS and reacted with 20 μM TCO-biotin for 30 min at 25 °C. The excess reagents were removed by methanol precipitation and the labeling intensity was analyzed by Western blot. For the labeling of BP, the reaction was used the same protocol but omitted the click reaction. For the labeling of AP, the click step was replaced by reacted with 100 μM azide-biotin, premixed 2-(4-((bis((1-tertbutyl-1H-1,2,3-triazol-4-yl)methyl)amino)methyl)-1H-1,2,3-triazol1-yl)-acetic acid (BTTAA)-CuSO_4_ complex (500 μM CuSO4, BTTAA:CuSO4 with a 2:1 molar ratio) and 2.5 mM freshly prepared sodium ascorbate for 2 h at 25 °C.

For the labeling of Hi-APEX in cell lysates, HEK293T cells were lysed in PBS with sonication and lysates were clarified by centrifugation at 20,000 g for 10 min at 4 °C. Protein concentration in clarified lysates was estimated with Pierce BCA Protein Assay Kit (Thermo Fisher) and normalized to 2 mg/mL. Lysates were incubated with 2 µM recombinant APEX2 and 500 µM of BP for 30 minutes, 500 µM of AP for 30 minutes or 500 µM of TP for 5 minutes, followed by a 1-minute vehicle or 1 mM H₂O₂ treatment. The click steps were as described above.

For the labeling of genetically-encoded Hi-APEX in living cells, HEK293T cells stable expressing APEX2 fusion construct of interest incubated with TP in DMEM for 5 minutes. The reaction was quenched by replacing the DMEM with an equal volume of quenching solution (10 mM ascorbate, 5 mM Trolox in PBS). Cells were washed with quenching solution twice. Cells were washed twice with 1 mL ice-cold PBS, harvested by scraping, pelleted by centrifugation at 1000 r.p.m. for 3 min, and either processed immediately or flash frozen in liquid nitrogen and stored at −80 °C before further analysis.

For the pulse-chase labeling of secretory proteins by Hi-APEX, HEK293T cells stably expressing APEX2-KDEL were incubated with 250 μM TP in DMEM for 5 min. Cells were washed trice with PBS. The buffer was changed to 10 mL fresh FBS-free DMEM and the cells were further cultured for 16 hours. Collect cells and culture media separately. The culture medium was centrifuged at 3000 rpm for 5 minutes, take the supernatant, and filtered through a 0.22 μm filter membrane for later use.

For the labeling of stress granule RNAs and proteins by Hi-APEX, HEK293T cells stably expressing G3BP1-APEX2 were treated with 500 μM sodium arsenite for an hour. At the last 5 minutes of sodium arsenite incubation, 500 μM TP was added for labeling. The reaction was quenched by replacing the DMEM with an equal volume of quenching solution (10 mM ascorbate, 5 mM Trolox in PBS). Cells were washed with quenching solution twice. Cells were washed twice with 1 mL ice-cold PBS, harvested by scraping, pelleted by centrifugation at 1000 r.p.m. for 3 min, and either processed immediately or flash frozen in liquid nitrogen and stored at −80 °C before further analysis.

For all cell labeling experiments described above, the cells were lysed by resuspending in RIPA lysis buffer (50 mM Tris pH 8, 150 mM NaCl, 0.1% SDS, 0.5% sodium deoxycholate, 1% Triton X-100, 1× protease inhibitor cocktail (Sigma-Aldrich)) by gentle pipetting and incubating for 5 min at 4 °C. Lysates were clarified by centrifugation at 20,000 g for 10 min at 4 °C. Protein concentration in clarified lysates was estimated with Pierce BCA Protein Assay Kit (Thermo Fisher) and normalized to 2 mg/mL. 50 μL of lysates were reacted with 20 μM TCO-biotin for 30 min at 25 °C. The excess reagents were removed by methanol and chloroform (4:1) precipitation and the labeling intensity was analyzed by Western blot.

For the pulse-chase labeling experiment, the culture medium was added to a 10 kDa ultrafiltration tube, centrifuge at 4000 rpm at 4 °C for 10 minutes, the centrifugation was repeated to concentrate all the culture medium. Add 3 mL of PBS, centrifuge at 4000 rpm at 4 °C for 10 minutes, repeat this process 3 times. After adding 1 mL of 1% SDS RIPA and soaking for about 1 hour, centrifuge at 4000 rpm and 4 °C for 10 minutes, and then the protein was harvested, quantified and analyzed as described above.

### Western blotting

For all Western blots, samples were resolved on a 10% SDS-PAGE gel. After SDS-PAGE, the gels were transferred to a PVDF membrane, and then stained by Ponceau S (5 min in 0.1% (w/v) Ponceau S in 5% acetic acid/water). The blots were then blocked in 5% (w/v) skim milk in TBS-T (Tris-buffered saline, 0.1% Tween 20) for at least 30 min at room temperature. For streptavidin blotting, the blots were stained with 0.3 μg/mL streptavidin-HRP (1:10,000 dilution; Beyotime, A0303) in TBS-T for 1 h at room temperature. For protein blotting, the blots were stained with ether anti-v5 (1:3,000 dilution, catalog no. 460705, Thermo Fisher) anti-FLAG(1:3,000 dilution, AE092, ABclonal) anti-GAPDH (1:3,000 dilution, AC001, ABclonal) anti-Actin (1:3,000 dilution, AC026, ABclonal) anti-TRMT61B (1:3000 dilution, 26009-1-AP, Proteintech) for 2 h at room temperature. After washing three times with TBS-T for 5 min each, the blots were stained with secondary antibodies in 5% BSA (w/v) in TBS-T for 1 h in room temperature. The blots were washed three times with TBS-T for 5 min each time before to development with Clarity Western ECL Blotting Substrates (Thermo Fisher) and imaging on the Tanon 5200multi.

### Confocal fluorescence microscopy

To validate the labeling of Hi-APEX at different organelle, HeLa cells were plated on 15-mm glass coverslips and transfected with APEX2 fusion construct of interest for 24 h. Cells were washed twice with PBS, and then incubated with 250 μM TP in DMEM for 5 min. The reaction was quenched by replacing the PBS with an equal volume of quenching solution (10 mM ascorbate, 5 mM Trolox in PBS). Cells were washed with quenching solution twice. Cells were washed once with 1 mL ice-cold PBS. Cells were fixed with cold methanol for 15 min at −20 °C and then washed three times with PBS. Tetrazine labeled proteins were then reacted with 100 μM TCO-biotin in PBS for 30 min at 25 °C. Cells were washed three times in PBS and blocked in 5% BSA dissolved in PBS for 1 h. The cells were incubated with primary antibodies against v5 (1:200 dilution) and a marker of each organelle including anti-citrate synthetase (1:200 dilution, catalog no. BM5202, BOSTER), anti-Calnexin (1:200 dilution, catalog no. A15631, ABclonal) overnight at 4 °C. Cells were washed three times in PBS with 5 min for each wash then incubated in secondary antibodies conjugated to either AlexaFluor-488 (1:200 dilution, catalog no. 33206ES60, Yeasen) or 568 (1:200 dilution, catalog no. ab175470, abcam) and streptavidin conjugated to AlexaFluor647 (1:300 dilution, catalog no. 35104ES60, Yeasen) for 1 h. Cells were washed five times in PBST (0.1% Tween-20 in PBS). Cells were incubated with DAPI (4′,6-diamidino-2-phenylindole, 1 μg/mL) for 10 min. After two additional PBS washes, samples were covered with aluminum foil and stored at 4 °C until imaging.

To validate the stress granule labeling by Hi-APEX, HEK293T cells stably expressing G3BP1-APEX2 fusion construct were plated on 15-mm glass coverslips and washed twice with PBS. The cells were treated with 500 μM sodium arsenite for an hour or not. At the last 5 minutes of sodium arsenite incubation, 500 μM TP was added for labeling. The reaction was quenched by replacing the DMEM with an equal volume of quenching solution (10 mM ascorbate, 5 mM Trolox in PBS). The remaining steps are as described above.

To validate the SG “orphan” proteins, HEK293T cells were treated with 500 μM sodium arsenite for an hour or not and washed with PBS for 3 times before fixed for 15 minutes with pre-cooled carbinol at −20 ℃. After 3 times washes in PBS, cells were incubated with primary antibodies against G3BP1(1:250 dilution, catalog no. A3968, ABclonal) and SPDL1(1:200 dilution, catalog no. 24689-1-AP, Proteintech) or CEP170(1:200 dilution, catalog no. 27325-1-AP, Proteintech) overnight at 4 °C. Cells were washed three times in PBS with 5 minutes for each wash then incubated in secondary antibodies conjugated to either AlexaFluor-488 and 568 for 1hour. Cells were washed 3 times in PBST (0.1% Tween-20 in PBS) and then incubated with DAPI (1:2000 dilution, catalog no. C1002, Beyotime) for 10 minutes. After two additional PBS washes, samples were covered with aluminum foil and stored at 4 °C until imaging.

To validate the neuron labeling of Hi-APEX, cultured neurons were infected with 10^9^ APEX2-mito AAV per well. After a week, the culture medium was changed to 500 μM TP in DMEM for 10 min for labeling. The reaction was quenched by replacing the DMEM with an equal volume of quenching solution (10 mM ascorbate, 5 mM Trolox in PBS). The remaining steps are as described above.

Confocal) microscope. Image processing and quantification were conducted using ImageJ2/FIJI software. Stress granule size was analyzed using the “Plot Profile” function to extract fluorescence intensity distributions along defined regions of interest. Stress granules (SGs) and cytosolic regions were segmented based on intensity thresholds, and quantitative measurements were performed on thresholded images acquired from randomly selected fields of view.

### Cell viability assay

For cytotoxicity assay, HEK293T cells stably expressing APEX2-mito cells or GPX4-overexpressing HT1080 cells were seeded at a density of 5000 cells per well in a 96-well plate overnight. Cell viability was quantified after being treated with TP or BP, or H_2_O_2_ for 3 minutes using MTS cell proliferation assay, or quantified after incubating with serum-free medium for 24 or 48 hours.

For RSL3-induced ferroptosis assay, HT1080 and other stably transfected cell lines were seeded at a density of 5,000 cells per well in a 96-well plate overnight. Then the cells were incubated with serum-free medium containing different concentrations of RSL3 for 24 hours. After washing with PBS, cell viability was measured using MTS cell proliferation assay (Promega).

### Filter-Aided Sample Preparation

For BSA labeling sites analysis, Filter-Aided Sample Preparation (FASP) was performed as previously described^58^. 50 μg of Hi-APEX *in vitro* labeled BSA was transferred to the 10K filter and centrifuged at 14,000 × g three times to remove the probe and finally diluted with 150 μL of PBS containing 8 M urea. 10 mM dithiothreitol (DTT) was added, and the mixture was incubated at 35°C for 30 minutes. Then, 20 mM iodoacetamide (IAA) was added, and the sample was incubated for another 30 minutes at 37°C in the dark. A 10K filter (Millipore) was equilibrated with 300 μL of 10 mM ammonium bicarbonate solution (ABC, pH 10). Each sample was centrifuged at 14,000 × g for 3 minutes at 16°C. An additional 200 μL of ABC solution was added to the 10K filter, and the centrifugation and addition cycle was repeated five times. 2 μg of trypsin was added to the proteome sample. The proteome samples in the 10K filter were digested for 16 hours at 37°C. After digestion, the samples were centrifuged at 20,000 × g for 10 minutes. The remaining material on the filter was washed twice by adding 200 μL of ABC solution and centrifuging at 20,000 × g for 10 minutes. The combined extracts were dried in a vacuum concentrator. The resulting peptide mixtures were reconstituted in 0.1% (v/v) formic acid (FA) in water prior to mass spectrometry analysis.

### Streptavidin enrichment

For the protein samples, cells in a 10 cm dish were lysed in 1 mL EDTA-free RIPA lysis buffer on ice for 30 minutes. Samples were centrifuged at 12,000g for 15 minutes at 4 ℃ and the protein concentration was normalized to a final protein concentration of 2 mg/mL using BCA Protein Assay Kit. The Supernatant was removed and subjected to click reaction with TCO-Biotin via IEDDA for 30 min at room temperature. After the click reaction, the proteins were extracted using chloroform-methanol precipitation and dissolved in RIPA buffer. 100 μL streptavidin magnetic beads (ChomiX Biotech Co., Ltd. Nanjing, China, cat. no. 02030002) for each sample were washed with RIPA twice and co-incubated with samples on a rotary shaker overnight at 4 ℃. Pellet the beads on a magnetic rack and wash the beads twice with 1 mL RIPA buffer, once with 1 mL of 1 M KCl, once with 1 mL of 0.1 M Na_2_CO_3_, once with 1 mL of 2 M urea in 10 mM Tris-HCl (pH 8.0), and twice with RIPA buffer. The enriched proteins with magnetic beads were resuspended in 500 μL 25 mM Ammonium bicarbonate (ABC) buffer with 6 M urea and 10 mM dithiothreitol (DTT) at 35 °C for 30 minutes and alkylated by addition of 20 mM iodoacetamide (IAA) at 37 °C for 30 minutes in the dark. The beads were then washed 6 times with 1 mL of 25 mM ABC buffer and resuspended in 200 μL of 25 mM ABC buffer and 10 ng/μL trypsin (Meizhiyuan). Trypsin digestion was performed at 37 °C on a thermomixer overnight. Collect the digested peptides in fresh microcentrifuge tubes, wash the beads once with 100 μL of 25 mM ABC buffer and once with 100 μL of 0.2% formic acid, the supernatant was combined with the peptide solution. The combined extracts were dried in a vacuum concentrator. The resulting peptide mixtures were reconstituted in 0.1% (v/v) formic acid (FA) in water prior to mass spectrometry analysis.

For the RNA samples, cells in a 10cm dish were lysed with 2mL TRIzol reagent. The homogenized sample was mixed and incubated with chloroform, the upper aqueous phase was pipetted out and subjected to RNA precipitation by adding 100% isopropanol. The RNA pellet was washed with 75% ethanol and dissolved in RNase-free water. The purified total RNA was then digested by DNase I (NEB, M0303S) at 37 °C for 30 min. The mixture was incubated with 100 μM TCO-biotin on a shaker at room temperature for 10 min. After click reaction, RNA was purified again with RNA Clean & Concentrator kit (Zymo, R1018, size limits were 17 nt to 23 kilobases) and was eluted into nuclease-free water. Then, 1 μg purified total RNA was taken out and set aside as INPUT. The remaining 100μg RNA was for enrichment.40 μl Dynabeads MyOne Streptavidin C1 (Invitrogen, 65002)beads for each sample were washed twice with 100 μl Binding & Washing(BW) buffer (100 mM Tris, pH 7.5, 1 M NaCl, 10 mM EDTA, 0.2% v/v Tween-20), twice with buffer A (0.1 M NaOH, 50 mM NaCl), twice with buffer B (100 mM NaCl) and finally re-suspended in 100 μl blocking buffer (1 μg μl^−1^ BSA, 1 μg μl^−1^ yeast tRNA in nuclease-free water) and incubated on a rotating mixer at room temperature for 2 h. The blocked beads were washed twice with BW buffer, and then mixed with purified RNA in 100 μl loading buffer. The mixture was incubated at room temperature for 45 min on a shaking incubator at 1,000 r.p.m. After removing the supernatant, the beads were washed with BW buffer three times, Urea buffer(4M Urea, 0.1%SDS in PBS) twice and PBS twice at room temperature. The RNA with beads was set aside for the following analysis as ENRICH.

### LC-MS/MS

For LC-MS/MS analysis, a loading column (100 µm × 2 cm) and a C18 capillary column (100 µm × 15 cm) packed in-house with Luna 3 µm C18(2) bulk material (Phenomenex, USA) were used to separate peptides. Separation was performed on a Vanquish™ Neo UHPLC system (Thermo Fisher Scientific) with mobile phase A consisting of water with 0.1% formic acid and mobile phase B composed of 94% acetonitrile with 0.1% formic acid. For protein samples, the LC gradient was initially held at 4% B for 1.8 min, then gradually increased to 5% at the 2-min mark, followed by a steady rise to 20% over the next 52 min, reaching 35% at 78 min and finally ramping up to 99% by 81.5 min. For superTOP-ABPP samples, the gradient remained at 4% B for the first 4 min before increasing to 20% over the next 105 min, then reaching 35% at 150 min and ultimately 99% at 159 min.

For samples analyzed on Q Exactive-Plus series Orbitrap mass spectrometers (Thermo Fisher Scientific), ionization was performed using an EASY-Spray source (Thermo Fisher Scientific) at +2.0 kV relative to ground, with an inlet capillary temperature maintained at 320 °C. Peptide precursor survey scans were acquired in the Orbitrap over a mass range of 350–1800 Th, with an AGC target of 300,000, a maximum injection time of 45 ms, and a resolution of 70,000 at 120 m/z. Monoisotopic precursor selection was enabled for peptide isotopic distributions, while precursors of z = 2–7 were selected for data-dependent MS/MS scans. Dynamic exclusion was applied for 25 s, with a ± 10 ppm window around the precursor monoisotope.

### Proteomic data analysis

The raw data were processed using DIA-NN (v2.2.0) in an advanced library-free module. The main search settings for in silico library generation were set as following: trypsin/P with maximum 3 missed cleavage; protein N-terminal M excision on; carbamidomethyl on C as fixed modification; oxidation on M as variable modification; peptide length from 5-30; precursor charge 1-4; precursor m/z from 250 to 1800; fragment m/z from 200 to 1800. The Human UniProt isoform sequence database (3AUP000005640) was used to annotate proteins for human cell samples. Other search parameters were set as following: quantification strategy was set to “QuantUMS (high precision)” mode; cross-run normalization was off; MS2 and MS1 mass accuracies were set to 0, allowing the DIA-NN to automatically determine mass tolerances; Scan window was set to 0 corresponding to the approximate average number of data points per peak; Peptidoforms and MBR were turned on; neural network classifier was single-pass mode.

For the tumor xenograft data filtering, all of the protein IDs whose missing value >3 were removed. The missing value imputation was conducted via the R package “imputeLCMD”(v2.1) with function “impute.MinPro”. The differential analysis was conducted using R package “DeqMS” (v1.26.0).

### RT–qPCR analysis of enriched RNA

For each sample, 1ug INPUT RNA and ENRICH RNA from 20 ug pre-enriched total RNA were reverse transcribed with random primers and ProtoScript II (NEB, E6560L) in 20 μl reaction buffer, according to the manufacturer’s instructions. Briefly, template RNA, random primer, ProtoScript II enzyme mix and reaction mix were mixed and incubated at 25 °C for 5 min, 42 °C for 1 h and 80 °C for 5 min. The INPUT and ENRICH cDNAs were aliquoted into four tubes (for four genes) as templates for qPCR. The templates were mixed with PowerUp SYBR Green Master Mix (Life, A25742) and primers, and then quantified by Vazyme Real-Time Quantitative Thermal Cycler FMR3 system. Ct values were averaged from three replicate measurements. Negative controls were treated in the same manner as the sample.

For each target, both forward and reverse primer sequences are listed. Oligonucleotide sequences are from 5’-end to 3’-end.

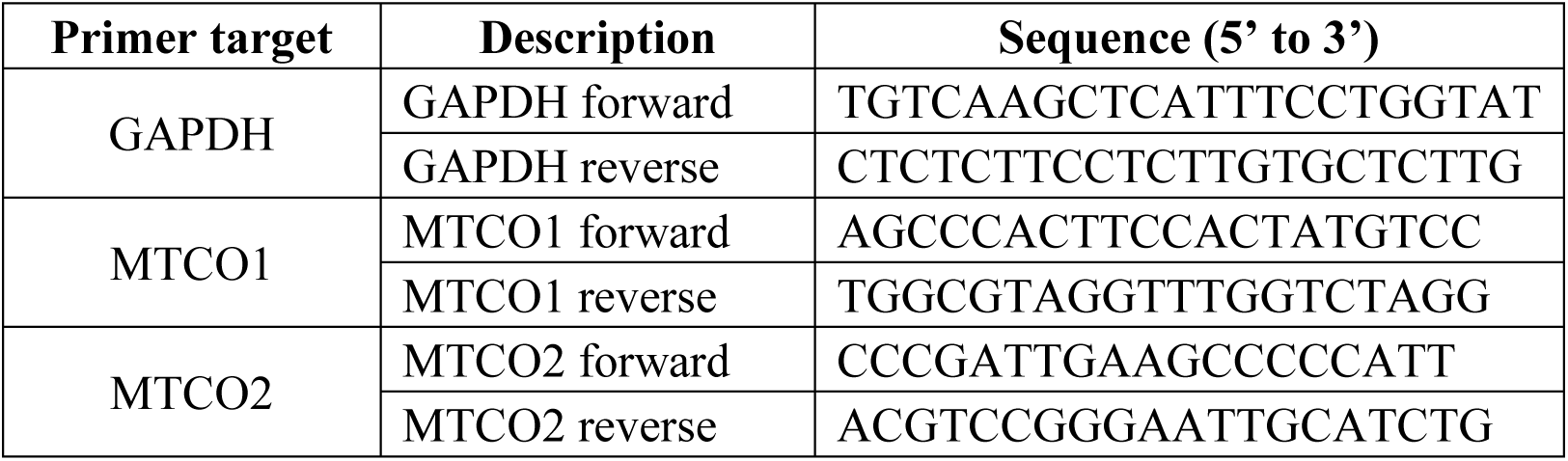
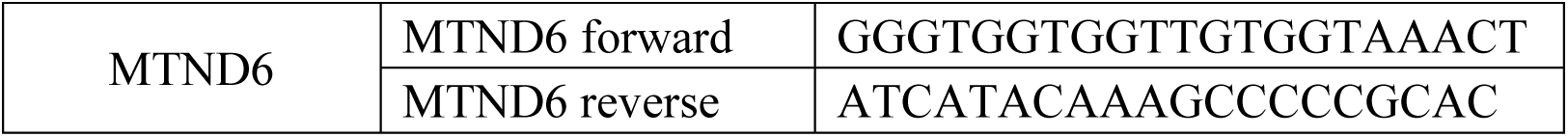

### Library construction for next-generation sequencing

VAHTS Universal V10 RNA-seq Library Prep Kit for Illumina (Vazyme, NR606) was used for Next-Generation Sequencing library construction. Then, 100 ng INPUT RNA or ENRICH RNA with beads from 100 ug pre-enriched total RNA was fragmentated in Frag/Prime Buffer at 85°C for 6 min and then reverse transcribed in the mixture with 7 μl of First Strand Buffer and 2 μl of First Strand Synthesis Enzyme Mix, following incubation at 25 °C for 10 min, 42 °C for 15 min, 70 °C for 15 min. Next, 25 μL of Second Strand Buffer with dUTP, 15 μl of Second Strand Enzyme Super Mix were added to the first strand reaction product for second strand cDNA synthesis and incubated for 30 min at 16 °C. For adaptor ligation, 5 μl 2uM Adaptor (Vazyme, N321), 25 μl Rapid Ligation Buffer and 5μl Rapid DNA Ligase were added to the 65 μl second strand cDNA and incubated for 15 min at 20 °C. Double-stranded cDNA was purified from the reaction mixture using VAHTS DNA Clean Beads (Vazyme, N411) by 0.6× beads binding, two rounds of 80% ethanol wash and 22 μl 0.1× TE buffer elution and then sorted using 0.6×/0.1× VAHTS DNA Clean Beads. Adaptor ligated DNA was amplified by PCR, and purified again with 0.9 × VAHTS DNA Clean Beads. During PCR, 25 μl VAHTS HiFi Amplification Mix, 5 μl primer mix and 20 μl adaptor ligated DNA were mixed and placed on a thermocycler with the following cycling conditions: 98 °C 45 s for one cycle, 98 °C 15 s with 60 °C 30s and 72°C 30s for 11 (INPUT samples) or 13(ENRICH samples) cycles, 72°C 1 min for one cycle, held at 4 °C. Every constructed libraries were equally mixed to 2 ng μl^−1^ and sent for high-throughput sequencing.

### Next-generation sequencing and data analysis

All sequencing data analyzed during this study are included in the Source Data. All cDNA libraries were deep-sequenced for 150 base paired reads on the Novaseq Xplus-PE150 platform(Novogene). Raw paired-end RNA-seq reads were subjected to adaptor and quality trimming using Trim Galore (v0.6.10). Reads shorter than 36 bp after trimming were discarded. Clean reads were subsequently aligned to hg38 reference genome (UCSC) using STAR (v2.5.2b) with default settings, allowing only uniquely mapped reads and excluding noncanonical splice junctions. Read counts per gene were calculated by featureCounts (v2.0.2), with strand-specific and paired-end counting enabled. The resulting raw count matrix was used for subsequent transcriptomic analyses.

To analyze the mitochondrial matrix Hi-APEX-seq data, we calculated FCs of genes with input count > 100. The genes with log2 (enrichment versus control) > 2 were identified as the enriched in the mitochondrial matrix dataset To analyze stress granule Hi-APEX-seq data, differential expression analysis was carried out by DESeq2^59^ (v.1.16.1). FCs and P values for each gene across different label conditions were reported.

RNA-seq BAM files from input and enrich groups (n=3 per group) were visualized in IGV (Broad Institute). Three genes (GAPDH, G3BP1, DCP2) were displayed, and coverage tracks were exported as SVGs for assembly in Adobe Illustrator. For mitochondrial transcripts, read coverage (log₂[reads + 1]) was computed in 50 bp bins and plotted with ggplot2, showing mean coverage per group.

### Co-immunoprecipitation

To validate the proximal proteins of mGPx4 identified by Hi-APEX, we performed co-immunoprecipitation assays using HT1080 cells stably expressing mGPX4-Flag. Briefly, cells grown in a 10 cm dish to approximately 80% confluency were treated with 150 nM RSL3 for 3 hours. Following treatment, the cells were washed three times with PBS, harvested by centrifugation at 1000 × g for 5 min, and resuspended in 600 µL of IP buffer (20 mM Tris-HCl, pH 7.5, 150 mM NaCl, 1% Triton X-100, 10% glycerol, 2.5 mM MgCl₂, and 0.5 mM CaCl₂) supplemented with 1× protease inhibitor cocktail and 20 U/mL DNase I. Cell lysis was conducted by rotating the suspension at 4 °C for 1 h to ensure complete disruption. The lysate was then centrifuged at 20,000 × g for 10 min at 4 °C to remove cellular debris. A 50 µL aliquot of the supernatant was mixed with SDS-PAGE loading buffer and boiled at 95 °C for 10 min to serve as the input sample. The remaining supernatant was incubated with pre-washed anti-FLAG M2 affinity gel beads (Sigma-Aldrich) at 4 °C overnight with gentle rotation. After incubation, the beads were washed three times with IP buffer, resuspended in 60 µL of 1× SDS-PAGE loading buffer, and boiled for subsequent Western blot analysis with the indicated antibodies.

### Animals

For xenograft and intravenous mouse model, male BALB/c nude mice aged 3–4 weeks and female C57BL/6J mice aged 6–7 weeks were obtained from Beijing Vital River Laboratory Animal Technology Co., Ltd (Beijing, China). The animals were housed in pressurized, individually ventilated cages (PIV/IVC) and maintained under specific-pathogen-free conditions, with free access to food and water in a controlled 12-h light/dark cycle. All animal experiments were conducted in accordance with the ethical standards and approved by the Institutional Animal Care and Use Committees of Tsinghua University (Beijing, China). The maximal tumor size/burden permitted by the Animal Care and Use Committee of Tsinghua University was 20 mm in diameter. We confirm that this limit was not exceeded during the course of the study.

For intracranial surgery experiment, female C57BL/6 J mice aged 7–8 weeks were obtained from the Laboratory Animal Center of the Institute of Genetics and Developmental Biology, Chinese Academy of Sciences (Beijing, China). The animals were housed in pressurized, individually ventilated cages (PIV/IVC) and maintained under specific-pathogen-free conditions, with free access to food and water in a controlled 12-h light/dark cycle. All animal experiments were conducted in accordance with the ethical standards and approved by the Animal Care and Use Committee of the Institute of Genetics and Developmental Biology, Chinese Academy of Sciences (Beijing, China).

### Xenograft Mouse Model

W.T. or stably expressing mGPx4-APEX2 HT1080 cells were resuspended in PBS with the density of 1 × 10^7^ cells per mL, and were injected into the flank of hind legs of BALB/c nude mice in a volume of 0.1 mL on both sides through s.c.. 2 weeks later, the tumor volume was about 100 mm^3^ measured by caliper, then mice were anesthetized and TP (5 mM solution in PBS with 5% Tween-80) was multi-point injected intratumorally in a total volume of 0.1 mL into both sides tumor. After 24h, repeat previous injection and mice were euthanized and tumors were removed in 1h after the second injection. The maximum tumor size approved by IACUC protocol was 2 cm in diameter and this was not exceeded in all experiments.

### Mice intracranial surgery

Adult mice were anesthetized with isoflurane inhalation, placed in a stereotaxic frame (RWD life science), and then using a Nanoliter 2000 microinjector (WPI), AAV vectors were delivered into the hippocampus at stereotaxic coordinates AP −1.8 mm from Bregma; ML ±1.3 mm; DV −1.6 mm from the dura. After infusion of the regents, the syringe needle was kept in place for 10 min to minimize the backflow of the regents. After 2 weeks of virus expression in mice brains, labeling experiments were performed.

For labeling in acute slice, mice were sacrificed. Brains were rapidly excised and submerged in the ice cold, oxygenated artificial cerebrospinal fluid (ACSF) composed of (in mM): 110 choline chloride, 2.5 KCl, 0.1 CaCl_2_, 3 MgCl_2_, 1.25 NaH2PO4, 25 NaHCO3, 3 Na pyruvate and 3 Inositol, then sectioned into 300 µm slices using a Leica VT1200 vibratome. Slices were incubated for at least 1h at 34 °C in oxygenated ACSF with 500μM TP.

For labeling in vivo, mice infected with the AAV were subjected to intracranial surgery again. For the injection, 500 nL PBS with TP (1 mM) was administered into the left and right hippocampus, respectively. Labeling was performed for 1 hours. After treatment, both experimental and control groups of mice were euthanized for tissue collection. For protein extraction, the mice brain was dissected out after euthanasia. The right and left hippocampal regions of the labeled site were isolated respectively and filled into fresh EP tubes. Samples were homogenized and lysed in RIPA. A small amount of sample was taken for western blot assay, the rest lysates were subjected to streptavidin purification following the procedure described above.

For histology, the mice were perfused with PBS and 4% PFA in sequence. Brains were dissected and post-fixed in 4% PFA overnight at 4 °C, and then transferred to 30% sucrose in PBS overnight at 4 °C. Dehydrated brain tissues were then embedded in O.C.T. compound (Tissue-Tek) and frozen on dry ice. Serial sections were coronally prepared at 15 μm for cryostat sections. For immunofluorescence staining, frozen brain sections were oven dried at 42 °C for 30 min, the brain slices were washed three times in PBS for 5 minutes each time. Then permeabilized in PBS with 0.3% Triton X-100 (PBST) for 20 min at room temperature, followed by washing three times in PBS for 5 minutes each time. Then incubate in PBS containing 2% normal goat serum for 1 hour at room temperature. Then incubated in 2% goat serum in PBS for 1 hour at room temperature. The slices were stained in 2% GS-PBST with anti-V5 (1:1000) antibodies at 4 °C for one overnight. Samples were washed three times in PBS at room temperature and incubated with anti-mouse-488, FITC-Streptavidin (1:1000) and DAPI (1:1000) in PBS for 1h, subsequently washed three times in PBS. The slices were mounted on glass slides in Fluoromount G (CITOTEST), followed with glass coverslips mounting on slides and sealing the edges with clear nail polish. The slides were stored at 4 °C overnight, images were acquired by Nikon A1 N-SIM S.

### Statistics and reproducibility

For STRING analysis, we selected a medium evidence level for the edges to represent the network. The evidence used to predict mGPx4 associations include protein homology, experimental evidence, text mining evidence, database evidence and co-expression evidence. Three replicates were performed for all experiments with similar results. For comparison between two groups, p values were determined using two-tailed Student’s t-tests, * p < 0.05; ** p < 0.01; *** p < 0.001; N.S. not significant. Error bars represent means ± SD. For GO analysis, we used DAVID bioinformatics for online analysis and the human proteome as background, the Benjamini–Hochberg method was performed to correct the t-tests.

## Notes

### Competing Interest Statement

The authors have declared no competing interest.

