## Supplementary information for "*In vivo*-compatible spatial multi-omics via hydrogen peroxide-independent APEX2 labeling"

[1]The State Key Laboratory of Membrane Biology, Tsinghua University, Beijing, 100084, China. [2]School of Pharmaceutical Sciences, Tsinghua University, Beijing, 100084, China. [3]Tsinghua-Peking Joint Center for Life Sciences, Tsinghua University, Beijing, 100084, China. [4]MOE Key Laboratory of Bioorganic Phosphorus Chemistry & Chemical Biology, Tsinghua University, Beijing, 100084, China. [5]Beijing Frontier Research Center for Biological Structure, Tsinghua University, Beijing, 100084, China. [6]Laboratory of Integrative Physiology, Institute of Genetics and Developmental Biology, Chinese Academy of Sciences, Beijing, 100101, China. [7]University of Chinese Academy of Sciences, Beijing, China, 101408, China. [8]State Key Laboratory of Female Fertility Promotion, Center for Reproductive Medicine, Department of Obstetrics and Gynecology, Peking University Third Hospital, Beijing, China. [9]Institute of Medicinal Plant Development, Chinese Academy of Medical Sciences & Peking Union Medical College, Beijing, China.

[#]These authors contribute equally.

### Supplementary figures

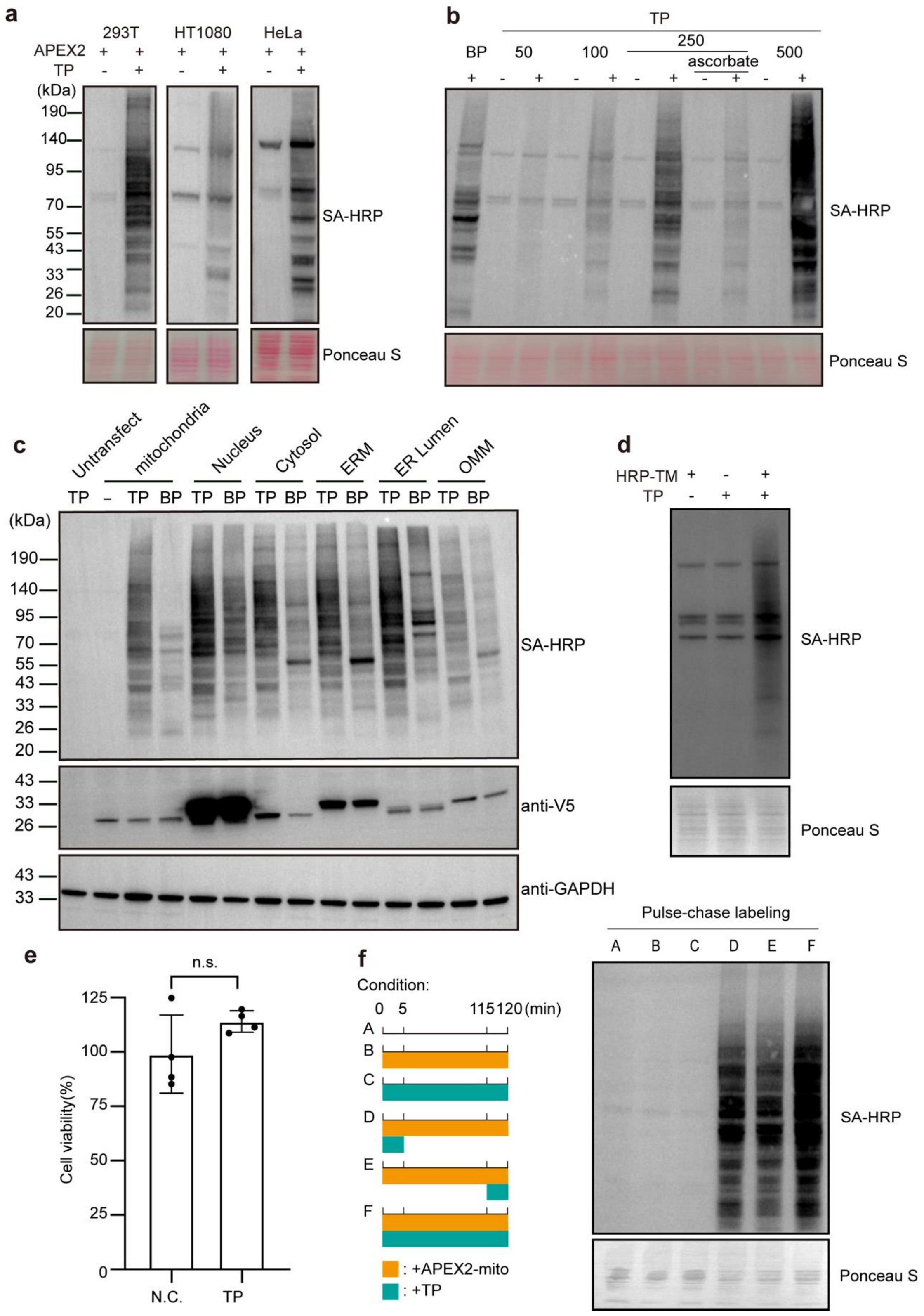


**Supplementary Fig. 1. Assessment of $H_2O_2$-independent APEX2 labeling using tetrazine-phenol (TP).**

(a) Mitochondrial matrix-localized APEX2 labeling with TP across multiple cell lines. HEK293T, HT1080, and HeLa cells expressing APEX2-mito were incubated with 200 μM TP for 30 min. After lysis, TP-labeled proteins were conjugated to TCO-biotin via an IDEEA reaction and detected by streptavidin blotting.

(b) Concentration dependence of APEX2 labeling with TP. HEK293T cells expressing APEX2-mito were treated with increasing concentrations of TP for 30 min. An additional sample treated with 250 μM TP was co-incubated with 10 mM sodium ascorbate.

(c) Comparison of TP-based APEX2 labeling across subcellular compartments. HEK293T cells expressing compartment-specific APEX2 constructs were treated with 200 μM TP for 30 min. For conventional biotin-phenol (BP) labeling, cells were incubated with 500 μM BP for 30 min, followed by 1 min stimulation with 1 mM $H_2O_2$.

(d) $H_2O_2$-independent labeling of HRP using TP. HEK293T cells expressing membrane-targeted HRP (HRP-TM) were incubated with 200 μM TP for 30 min.

(e) Cell viability assay following a 5 min treatment with 200 μM TP in APEX2-mito-expressing cells. n.s., not significant. Data are presented as mean ± SD.

(f) Pulse-chase analysis of APEX2 labeling with TP. APEX2-mito-expressing HEK293T cells were pulsed with 200 μM TP for 5 min, washed, and chased in vehicle medium for 115 min. Cells were then lysed and analyzed by click chemistry-based in-gel fluorescence. A schematic of the labeling strategy is shown on the left.

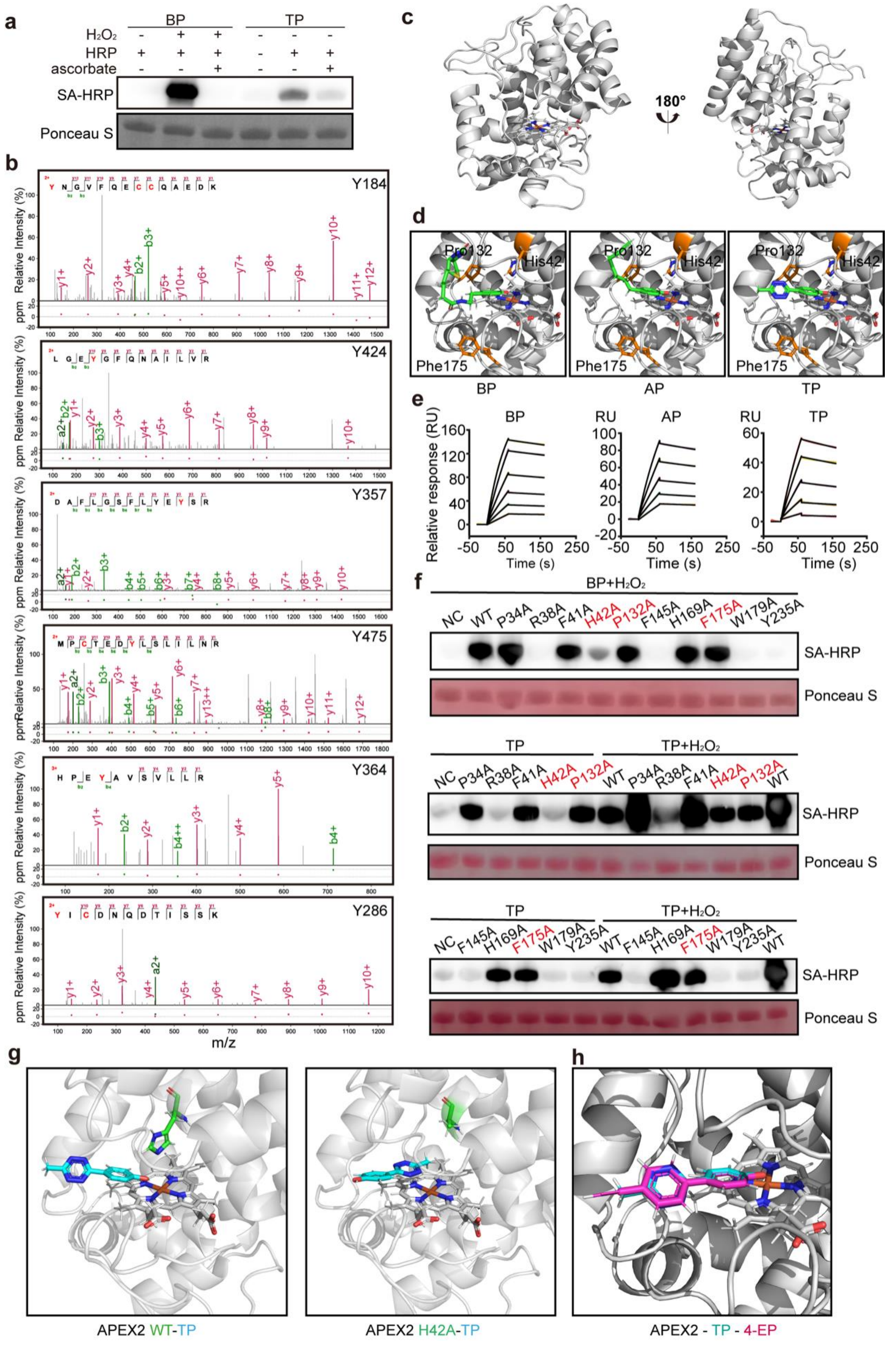


**Supplementary Fig. 2.** ***In vitro*** **APEX2 labeling with TP.**

(a) $H_2O_2$-independent labeling of BSA by recombinant HRP using TP. For TP labeling, a mixture of 2 μM HRP and 2 mg/mL BSA was incubated with 500 μM TP for 30 min. For BP labeling, the same HRP/BSA mixture was incubated with 500 μM BP for 30 min, followed by 1 min stimulation with 1 mM $H_2O_2$. To probe the labeling mechanism, each condition was also treated with 10 mM sodium ascorbate. TP-labeled proteins were conjugated to TCO-biotin via an IDEEA reaction, and all samples were analyzed by streptavidin blotting.
(b) Representative MS2 spectra of BSA peptides bearing TP labeling.
(c) Crystal structure of APEX2 (PDB: 9X9L).
(d) Molecular docking of BP, AP, and TP into the apo form of APEX2.
(e) Surface plasmon resonance (SPR) analysis of APEX2 binding interactions with BP, AP, and TP.
(f) Streptavidin blotting assessment of $H_2O_2$-independent TP labeling using APEX2 mutants.
(g) Molecular docking of TP into wild-type APEX2 and the H42A mutant.
(h) Molecular docking of 4-EP into wild-type APEX2.

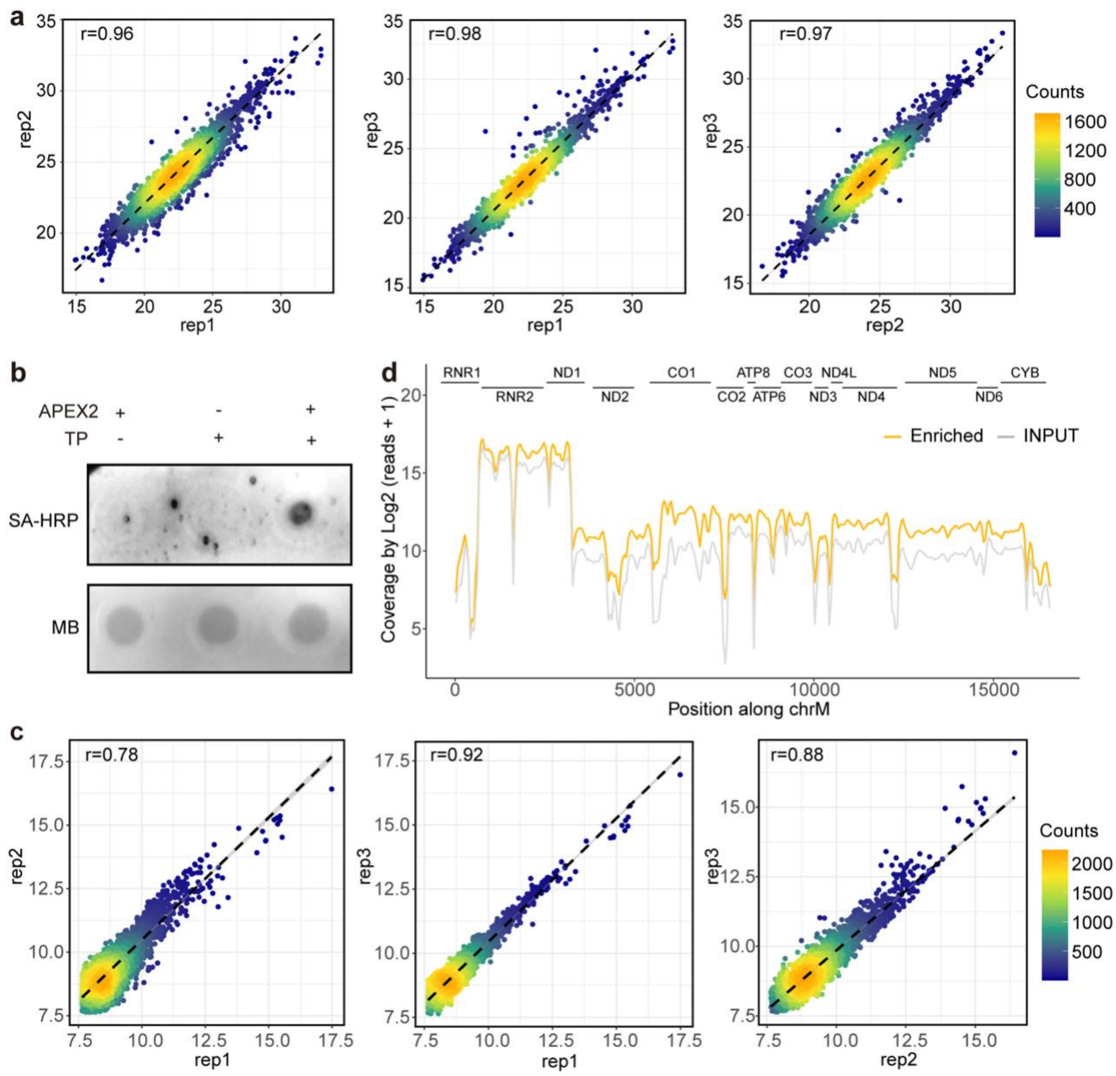


**Supplementary Fig. 3. Hi-APEX profiling of the mitochondrial matrix.**

(a) Correlation of protein enrichment ratios across biological replicates in mitochondrial matrix Hi-APEX proteomic profiling.

(b) Streptavidin dot blot analysis of total RNA following Hi-APEX mitochondrial labeling. HEK293T cells expressing APEX2-mito were treated with 500 μM TP for 5 min. Total RNA was extracted, click-conjugated to TCO-biotin, and analyzed by streptavidin dot blotting.

(c) Correlation of RNA enrichment ratios across biological replicates in mitochondrial matrix Hi-APEX transcriptomic profiling.

(d) Line plot showing mitochondrial genome coverage in the input and enriched fraction of Hi-APEX-mito labeled samples.

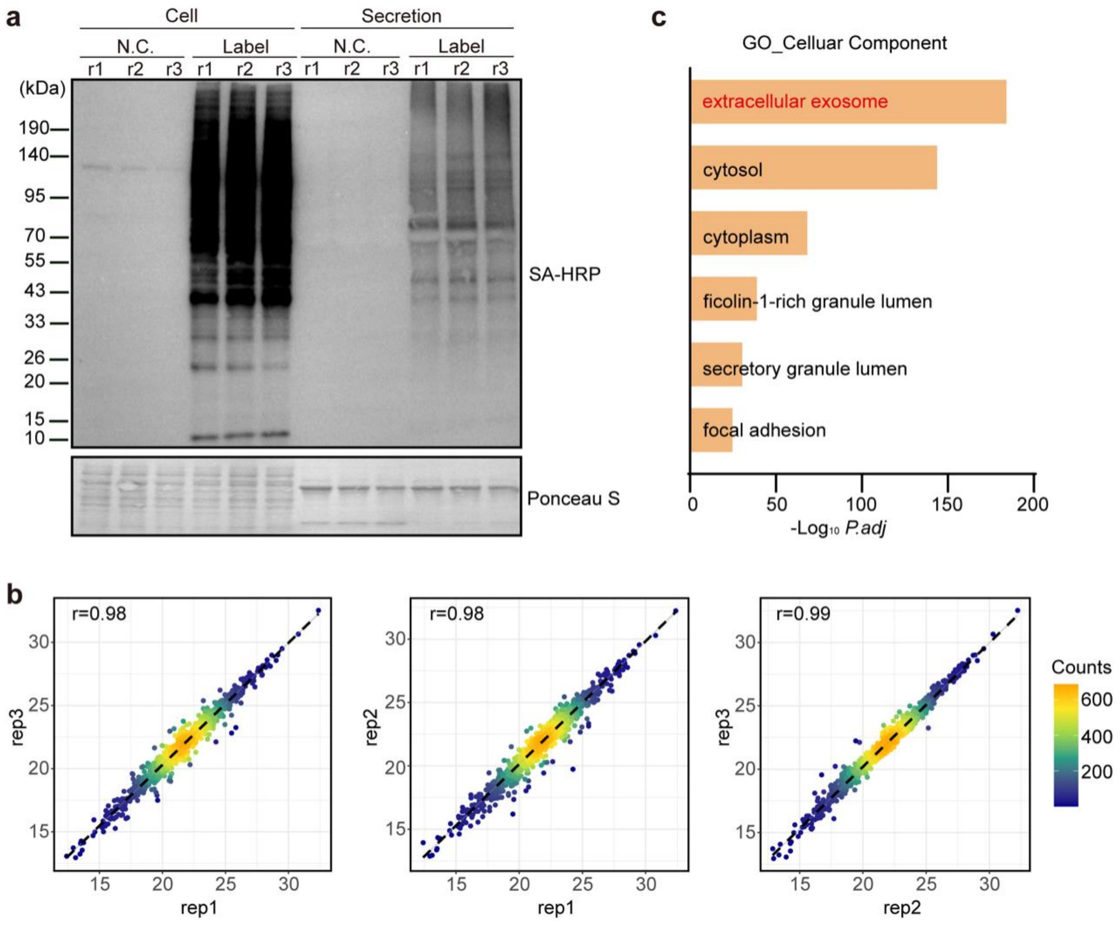


**Supplementary Fig. 4. Hi-APEX profiling of ER-derived secretomes.**

(a) Streptavidin blot analysis of cell lysates and conditioned medium following the pulse-chase labeling procedure outlined in Figure 3h.

(b) Correlation of protein enrichment ratios across biological replicates in the Hi-APEX secretome profiling.

(c) GOCC analysis of proteins enriched by Hi-APEX-KDEL pulse-chase labeling.

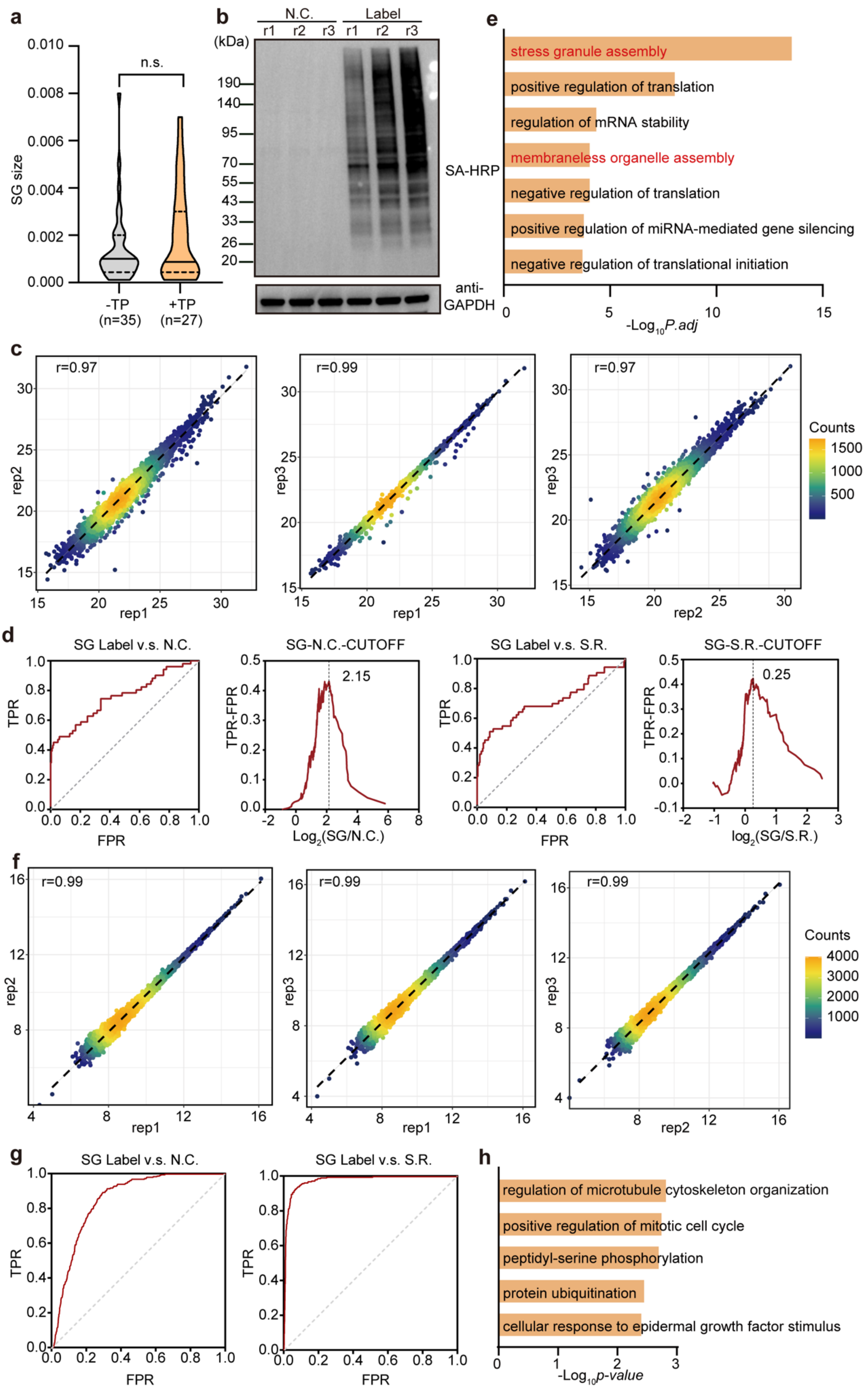


**Supplementary Fig. 5. Simultaneous proteomic and transcriptomic mapping of stress**

**granules using Hi-APEX.**

(a) Size distribution of stress granules with and without Hi-APEX-G3BP1 labeling. n.s., not significant.
(b) Representative streptavidin blot analysis of Hi-APEX-G3BP1 labeling efficiency.
(c) Correlation of protein enrichment ratios across biological replicates in Hi-APEX-G3BP1 proteomic profiling.
(d) ROC analysis of protein enrichment using negative control (N.C.) or spatial reference (S.R.) datasets in Hi-APEX-G3BP1 proteomics. True positives represent known stress granule proteins; false positives are nuclear proteins.
(e) GOCC analysis of proteins enriched by Hi-APEX-G3BP1 labeling.
(f) Correlation of RNA enrichment ratios across biological replicates in Hi-APEX-G3BP1 transcriptomic profiling.
(g) ROC analysis of RNA enrichment using negative control (N.C.) or spatial reference (S.R.) datasets in Hi-APEX-G3BP1 transcriptomics. True positives represent known SG-proximal transcripts; false positives are known SG-excluded transcripts.
(h) GOBP analysis of transcripts enriched by Hi-APEX-G3BP1 labeling.

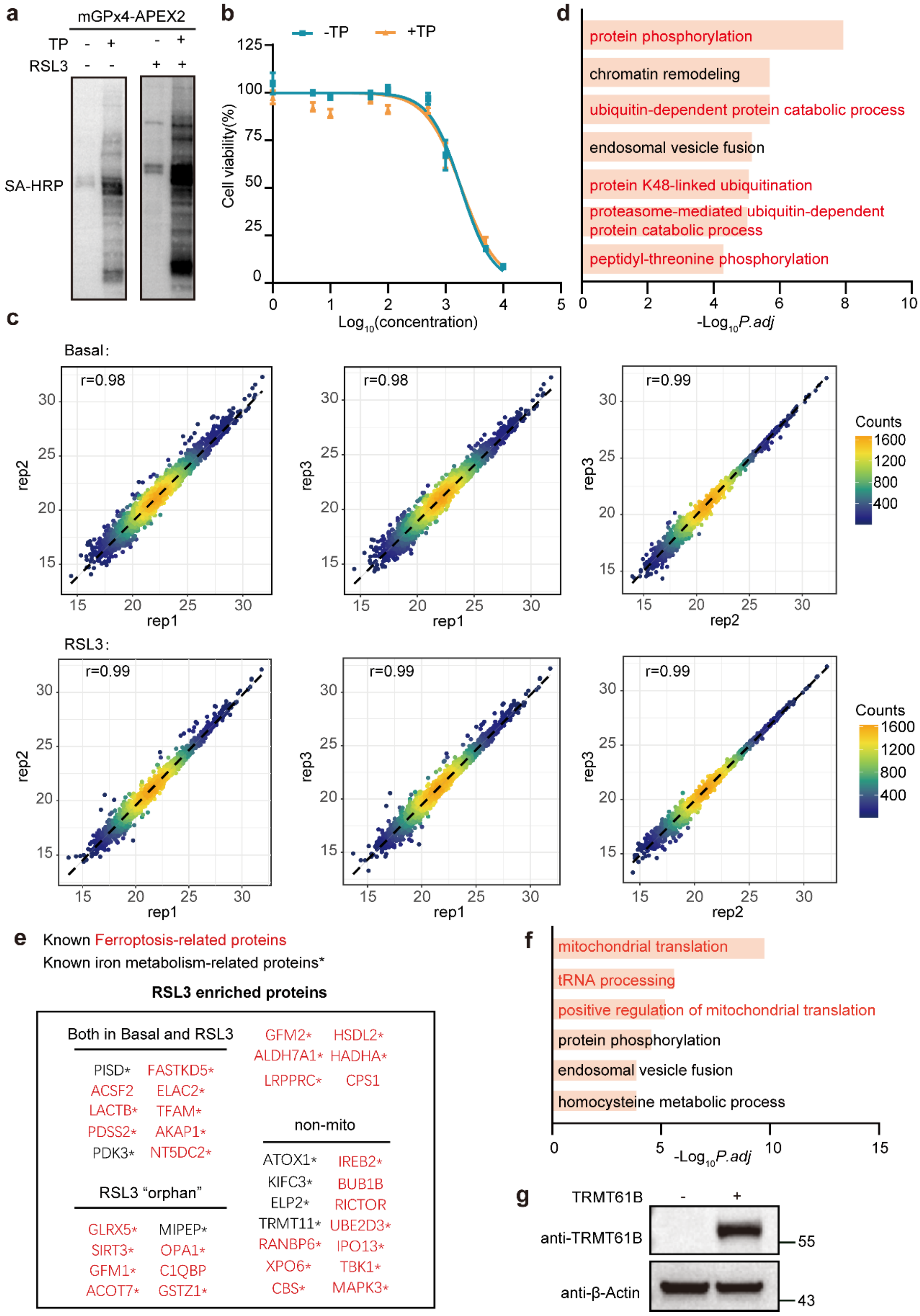


**Supplementary Fig. 6. Mapping the mitochondrial GPx4 interactome during ferroptosis using Hi-APEX.**

(a) Streptavidin blot analysis of Hi-APEX-labeled mGPx4-interacting proteins under basal and RSL3-treated conditions.

(b) Cell viability assay of mGPx4-APEX2 cells treated with 200 μM TP for 5 minutes. Data are presented as mean ± SD.
(c) Correlation of protein enrichment ratios across biological replicates in mGPx4-APEX2 proteomic profiling under basal and RSL3-treated conditions.
(d) GOBP analysis of non-mitochondrial proteins enriched by mGPx4-APEX2 profiling under RSL3 treatment.
(e) Ferroptosis- and iron metabolism-related proteins among the top-enriched interactors identified by Hi-APEX-mGPx4.
(f) GOBP analysis of all proteins enriched by mGPx4-APEX2 profiling under RSL3 treatment.
(g) Validation of TRMT61B overexpression.

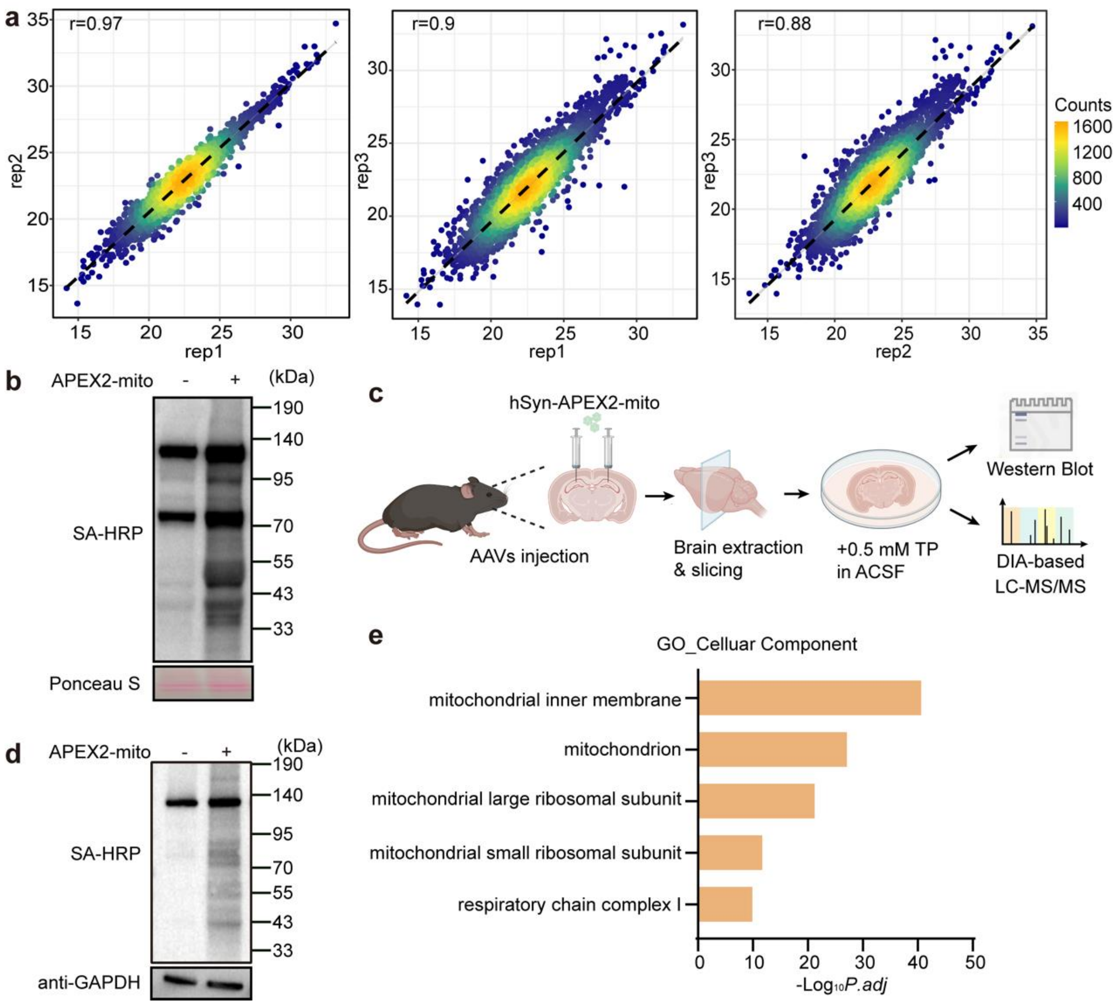


**Supplementary Fig. 7. *In vivo* Hi-APEX labeling.**

(a) Correlation of protein enrichment ratios across biological replicates in mGPx4-APEX2 proteomic profiling from tumor xenografts.

(b) Representative streptavidin blot analysis of mitochondrial matrix-targeted Hi-APEX labeling in cultured cortical neurons.

(c) Schematic of hippocampal neuronal mitochondrial matrix labeling using Hi-APEX. Neuron-specific AAVs encoding mitochondrial matrix-targeted APEX2 were injected into the hippocampus. After two weeks, brains were sectioned into 300 µm slices in ACSF, followed by incubation with 500 µM TP for 30 minutes.

(d) Representative streptavidin blot analysis of mitochondrial matrix-targeted Hi-APEX labeling in hippocampal brain slices.

(e) GOCC analysis of proteins enriched by mitochondrial matrix-targeted Hi-APEX labeling in brain slices.

**Supplementary tables**

**Table S1. Proteomic profiling of the mitochondrial matrix via Hi-APEX. Related to Figure 3 and Supplementary Fig 3.** The average ratio for each protein is shown. Proteins with an average Hi-APEX/control ratio greater than 2.72 and a $p$-value < 0.05 (one-sided t-test) were classified as enriched. The complete dataset of all identified and quantified proteins, including intensities, is provided in Tab 1. Enriched proteins are annotated in Tab 2, and the corresponding list is shown in Tab 3.

**Table S2. Transcriptomic profiling of the mitochondrial matrix via Hi-APEX. Related to Figure 3 and Supplementary Fig 3.** The counts for each labeled transcript are shown. Transcripts with an average Hi-APEX/control ratio greater than 2 and a $p$-value < 0.05 (one-sided t-test) were classified as enriched. The complete dataset of all identified and quantified transcripts, including intensities, is provided in Tab 1.

**Table S3. Proteomic profiling of the ER-originated secretion via Hi-APEX. Related to Figure 3 and Supplementary Fig 4.** The average ratio for each labeled protein is shown. Proteins with a Hi-APEX/control ratio greater than 10.27 and a $p$-value < 0.05 (one-sided t-test) were classified as enriched. The complete dataset is provided in Tab 1. Enriched proteins are annotated in Tab 1, and the corresponding list is shown in Tab 2.

**Table S4. Proteomic profiling of stress granules via Hi-APEX. Related to Figure 4 and Supplementary Fig 5.** The LABEL/negative control (N.C.) and LABEL/spatial reference (S.R.) ratios for each labeled protein are shown. Proteins with a LABEL/N.C. ratio > 4.44 and $p$-value < 0.05 were defined as enriched and are listed in Tab 1. Subsequently, proteins with a LABEL/S.R. ratio > 1.19 and $p$-value < 0.05 were defined as enriched and are listed in Tab 2. Proteins meeting both enrichment criteria were defined as *bona fide* stress granule proteins; these are listed and annotated in Tab 3, while the related protein list is shown in Tab 4. All $p$-values were determined by the one-sided t-test.

**Table S5. Transcriptomic profiling of stress granules via Hi-APEX. Related to Figure 4 and Supplementary Fig 5.** The LABEL/N.C. and LABEL/S.R. ratios for each labeled transcript are shown. Differential expression analysis was performed using the R package "DESeq2". Transcripts with a LABEL/N.C. ratio > 1.41 and an adjusted $p$-value < 0.05 were defined as enriched and are listed in Tab 1. Subsequently, transcripts with a LABEL/S.R.

ratio > 1.41 and *p*-value < 0.05 were defined as enriched, while those with a ratio < 0.71 and *p*-value < 0.05 were defined as depleted; both groups are listed in Tab 2. Based on these comparisons, transcripts enriched in both datasets were defined as stress granule-enriched transcripts, whereas transcripts enriched in the LABEL/N.C. comparison but depleted in the spatial reference were defined as stress granule-depleted transcripts. Both categories are listed and annotated in Tab 3, and the related transcript list is shown in Tab 4. *P*-values were determined by the one-sided t-test.

**Table S6. Proteomic profiling of mGPx4-proximal proteins in cultured cells under basal and ferroptosis conditions by Hi-APEX. Related to Figure 5 and Supplementary Fig 6.** In both basal and ferroptosis conditions, proteins with a Hi-APEX/control ratio > 2 and *p*-value < 0.05 (one-sided t-test) were classified as enriched. The complete dataset of all identified and quantified proteins is provided. Enriched proteins under basal and ferroptosis conditions are listed in Tabs 1 and 2, respectively. Proteins enriched during ferroptosis are annotated in Tab 3, and the related list is shown in Tab 4.

**Table S7. Proteomic profiling of mGPx4-proximal proteins in tumor xenografts by Hi-APEX. Related to Figure 6 and Supplementary Fig 7.** The average ratio for each labeled protein is shown. Proteins with an average Hi-APEX/control ratio greater than 2 and a *p*-value < 0.05 were classified as enriched. Differential analysis was performed using the R package "DEqMS". The complete dataset of all identified and quantified proteins is provided in Tab 1

**Table S8. Proteomic profiling of mitochondria proteins in mouse neuron *in vivo* by Hi-APEX. Related to Figure 6 and Supplementary Fig 7.** The average ratio for each labeled protein is shown. Proteins with an average Hi-APEX/control ratio greater than 2 and a *p*-value < 0.05 were classified as enriched. The complete dataset of all identified and quantified proteins of mouse hippocampus and brain slices is provided separately in Tab 1 and 2.
